# Expression and Engineering of Conductive Cytochrome Nanowires

**DOI:** 10.64898/2026.07.18.739370

**Authors:** Eric R. Szmuc, Xiaomeng Liu, Marcel Lopez Reed, Vidhika Damani, David Walker, Axel Brilot, Lizette Lozano-Zambrano, Guihua Yu, Benjamin K. Keitz, Andrew D. Ellington

## Abstract

Electrically conductive protein nanowires produced by metal-reducing bacteria have attracted interest as sustainable electronic materials, but their study and engineering have been limited by difficulties in expression, purification, and genetic manipulation. Here we establish *Shewanella oneidensis* as a heterologous host for production of *Geobacter sulfurreducens* OmcZ nanowires and progress a complementary *in vitro* assembly strategy that yields highly pure nanowire preparations. This platform enabled systematic engineering of OmcZ, revealing extensive tolerance to sequence variation and faciliating the design of enhanced-conductivity variants. Guided by comparative analysis of environmental OmcZ homologs, we generated a chimeric nanowire, OmcZ^+^, that exhibited a ∼3.5-fold increase in bulk conductivity while retaining the overall structure of the parent nanowire. Cryo-electron microscopy revealed unexpected architectural plasticity in OmcZ^+^ wires, including non-linear dendritic and pentameric assemblies mediated by the solvent-exposed heme VII, suggesting previously unrecognized modes of cytochrome nanowire organization. Incorporation of engineered OmcZ variants into water evaporation-induced electricity generators produced power densities up to ∼25.3 μW cm⁻² and enabled high-performance operation in saline environments, including seawater and human sweat. Together, these results establish a versatile platform for the production, structural analysis, and engineering of cytochrome nanowires, providing a foundation for the development of programmable biological electronic materials.

## INTRODUCTION

Multiheme cytochromes are promising materials for next-generation bioelectronics due to their combination of high conductivity and genetic programmability. Amongst the most promising class of multiheme cytochromes are those that assemble into nanowires, forming protein-based electronic conductors with exceptional charge-transport properties. For example, the octaheme cytochrome OmcZ from *Geobacter sulfurreducens* exhibits conductivities exceeding 30 S·cm⁻^1^, rivaling synthetic conductors such as PEDOT:PSS^2^, while maintaining the molecular programmability of a protein. High-resolution cryo-electron microscopy and electrochemical measurements of OmcZ wires have revealed a stacked heme array architecture where sub-6 Å heme spacing facilitates multistep electron hopping and charge delocalization along the filament axis^3^, ultimately enabling efficient charge transport and intrinsic redox activity.

In *G. sulfurreducens*, OmcZ nanowires form in the extracellular matrix through a biologically regulated assembly pathway in which the serine protease OzpA cleaves the pro-protein OmcZ_50_ into the active, nanowire forming OmcZ_303_. Once formed, OmcZ nanowires display remarkable stability, withstanding temperatures up to 98 °C and maintaining electronic performance for months under ambient conditions^1,3^, underscoring their potential as robust biological conductors. Despite its exceptional properties, OmcZ (both in monomeric and wire form) has been challenging to produce at scale due to strict expression requirements within *Geobacter* biofilms and concomitant difficulties with purification. For example, OmcZ expression primarily occurs in anoxic biofilm communities formed on the surface of high potential electrodes^4,5^ with planktonic cells exhibiting little to no OmcZ production^1^. While the heterologous expression of OmcZ in *Escherichia coli* has been demonstrated^3^, it is highly inefficient due to a lack of native cytochrome maturation machinery and the required co-expression of the roughly 6.5kb *ccmABCDEFGH* operon responsible for c-type cytochrome maturation^6^. With no native extracellular cytochrome expression pathway, *E. coli* also lacks a dedicated route to ensuring the translated apocytochrome is properly localized to the heterologously expressed cytochrome maturation machinery, fundamentally limiting apocytochrome heme incorporation and ultimately holocytochrome formation.

Given these limitations, we hypothesized that another model electroactive bacterium, *Shewanella oneidensis*, may prove a suitable host for OmcZ expression. In contrast to *E. coli*, *S. oneidensis* naturally encodes 42 putative c-type cytochromes^7^, which support robust extracellular electron transfer for coupling oxidative metabolism to the reduction of a wide range of metal oxides and inorganic substrates^8,9,10,11,12^. Similar to *Geobacter*, the extracellular electron transfer network in *S. oneidensis* has been extensively characterized^13,14^ and genetically engineered^15,16^, and numerous established cytochrome knockouts and expression variants are available that could potentially free metabolically expensive heme maturation resources for amplified synthesis of heterologous multiheme cytochromes.

Here, we demonstrate that *S. oneidensis* is an excellent host for OmcZ expression and subsequently optimize OmcZ nanowire production *in vivo* and *in vitro*, including identifying genetic variants of OmcZ with enhanced conductivity. Our studies also reveal a surprising structural plasticity during nanowire assembly, with some OmcZ variants forming pentameric rings and dendritic branching, suggesting intriguing opportunities for engineering novel electronic structures. We also leverage our ability to produce large amounts of OmcZ and enhanced variants to develop evaporation-induced power generation devices with outstanding performance. The large-scale synthesis of genetically programmable, electron-conducting nanowires should facilitate their integration into real-world bioelectronic power generating devices and provide a firm basis for establishing biologically derived electronic materials as a foundation for next-generation soft, self-powered, and environmentally benign devices.

## RESULTS

### OmcZ expression and nanowire formation in *Shewanella oneidensis*

To efficiently express *Geobacter* OmcZ as an extracellular, cytochrome-containing nanowire in *S. oneidensis* (WT strain MR-1), a three gene operon consisting of the pro-protein *omcZ_50_*, *ozpA*, and a prolyl isomerase was cloned into a pET expression plasmid (**Fig.1a**, **Table S1**). The entire operon was placed under the control of LacI, **Fig.1b**. Heterologous expression, periplasmic processing, and secretion were facilitated by replacing the N-terminal signal sequences of OmcZ_50_ and OzpA with *Shewanella*-specific alternatives. Since OmcZ expression in *Geobacter* normally occurs via the signal peptidase I (SPI) pathway the *Shewanella* SPI signal sequence from cctA, previously shown to secrete the endogenous cytochrome UndA1 into culture supernatant^17^, was added to the *Geobacter* OmcZ gene. Similarly, the signal sequence from the *Shewanella* sapSH extracellular subtilase (WP_011073002, 31% identity to *Geobacter* OzpA) was used to achieve extracellular OzpA expression. Two structural variants of OmcZ were initially screened, containing either the native, but atypical, motif of heme II (CX_14_CH) or in its place an engineered, canonical heme II motif (CXXCH; termed OmcZ[Δ56-67]; **Fig.S1**). While previous efforts in *E.coli*^18^ have suggested that native OmcZ expression should be possible, we were unable to demonstrate expression of OmcZ containing the atypical heme motif in *S. oneidensis*. Going forward, all experiments were carried out in *S. oneidensis* with OmcZ[Δ56-67] (now referred throughout as OmcZʹ). Unless otherwise indicated, all subsequent sequence substitutions and analyses refer to this OmcZʹ scaffold, while the native *G. sulfurreducens* protein is explicitly referred as WT OmcZ.

Upon IPTG induction during anaerobic growth, we measured an increase in absorbance at 410nm consistent with cytochrome production and export to the extracellular space (**Fig.1c**). Nanowires roughly 4nm in diameter were visible in transmission electron micrographs (TEM) taken of whole cells (**Fig.S2**). These wires exhibited similar angular structure to WT OmcZ and were distinct from the rod-shaped pili and flagellum typically formed by *S. oneidensis* (**Fig.S2**). Heme staining of gel-separated protein monomers^19^ revealed bands consistent with both OmcZʹ_50_ and OmcZʹ_30_ (**Fig.S3**), while samples lacking the OmcZʹ expression cassette showed no such bands, nor identifiable filaments with morphologies similar to OmcZ nanowires (**Fig.S2)**. Together, these results indicate that *S. oneidensis* is a suitable host for OmcZʹ expression *in vivo*.

### *In vitro* synthesis of OmcZʹ nanowires

Having demonstrated heterologous expression of OmcZʹ nanowires in *S. oneidensis*, we next attempted to synthesize pure wires from purified proteins *in vitro*. The three gene *omcZ* operon was divided into two plasmids: one encoding OmcZʹ_50_ and the prolyl isomerase, and the other encoding OzpA and the prolyl isomerase, an arrangement similar to Gu et al. (2023) in *E. coli* (**Fig.1e**). StrepII affinity tags were added to the C-terminus of OmcZʹ and to the N-terminus of OzpA to facilitate purification. After initial expression attempts failed for OzpA (**Fig.S4**) the sapSH signal sequence, which may dictate outer membrane association (**Fig.S5**), was replaced with the signal peptide from ompA (WP_330113719).

The parallel purification of OzpA and OmcZʹ from *Shewanella* supernatants yielded high protein titers: ∼2mg/L and ∼16mg/L, respectively. SDS-PAGE gel staining via Coomassie (OzpA) and heme-stain (OmcZʹ) confirmed the size and purity of both proteins (**Fig.1f**). The identity of OmcZʹ_50_ was further confirmed through LC/MS and mass spectroscopy (**Fig.1g, Fig.S6**), with LC/MS identifying 55% sequence coverage and mass spectroscopy identifying a size of 51.986 kDa, consistent with OmcZʹ_50_ containing 8 heme cofactors and the StrepII affinity tag. *In vitro* formation of OmcZʹ nanowires was initiated by incubating purified OmcZʹ_50_ with activated OzpA at room temperature. Gel electrophoresis and heme staining of OmcZʹ samples taken before and after OzpA incubation revealed a significant increase in the amount of OmcZʹ_30_ following incubation, with minimal uncleaved material remaining (**Fig.1h**). After 24 hours, extensive nanowire formation was observed in sample TEM images taken following incubation (**Fig.1i**). These data further confirm that OzpA is responsible for OmcZʹ_50_ pro-protein cleavage and subsequent OmcZʹ_30_ self-assembly into nanowires^3^, and that *S. oneidensis* serves as an excellent host for the *in vitro* production of highly pure OmcZʹ nanowires (**Fig.S7**).

### Expression optimization with OmcZʹ-GFP fusion

To further improve OmcZʹ expression in *S. oneidensis,* native cytochrome expression was manipulated to free up heme resources. Specifically, OmcZʹ expression was assessed in the cytochrome knockout strains *Δ3* (*ΔmtrCΔomcAΔmtrF,* JG596)^15^, *Δ8* (*ΔmtrAΔdmsEΔmtrDΔcctAΔSO_4360ΔmtrCΔmtrFΔOmcA,* JG1194)^16^, and *Δ10* (*ΔmtrBΔmtrCΔomcAΔmtrFΔmtrAΔmtrDΔdmsEΔSO_4360ΔcctAΔrecA,* JG1519)^16^. These strains retain *fccA* and *cymA*, which are necessary for fumarate reduction and planktonic, anaerobic growth in *S. oneidensis*^20^. Because absorbance may be confounded by varying genomically encoded cytochromes, we constructed a C-terminal OmcZʹ_50_-GFP fusion^21^ to more readily assay expression and fluorescence was monitored in the supernatant of an anaerobic cell culture after overnight maturation to facilitate GFP fluorophore formation (**Fig.2a**). As expected, cell density normalized fluorescence values increased only in the presence of the expression cassette and induction with IPTG. Only the cytochrome knockout *Δ3* outperformed wild-type *Shewanella* MR-1 for OmcZʹ expression, with a ∼120% improved yield, **Fig.2b**, while further knockouts slightly impaired OmcZʹ production. We also measured a strong correlation between GFP fluorescence and supernatant absorbance at 410nm (**Fig.S8**), confirming this assay is a suitable reporter of OmcZ expression.

Using our OmcZ’-GFP fusion, we next optimized media and growth conditions for OmcZ’ expression, **Fig.2c**. Intermediate IPTG concentrations resulted in superior OmcZ’ production while higher concentrations resulted in minimal protein expression. Since *S. oneidensis* cytochrome expression is affected by available iron^22^, we attempted iron supplementation and found that ≥30µM added iron had a beneficial impact on OmcZʹ expression, with supernatants exhibiting over 1.5X greater fluorescence response over media relying on yeast extract (∼1.7µM under conditions used) alone (**Fig.2d**). Finally, the OmcZʹ-GFP fusion was used to characterize the role of the prolyl isomerase in OmcZ expression. In the presence of a prolyl isomerase deletion (ΔPI; **Fig.2e**) fluorescence was reduced, indicating active participation in proper OmcZʹ folding and extracellular expression. Heme-straining in the absence of prolyl isomerase revealed significant reductions in band intensity (**Fig.S9**), further suggesting impairment of proper folding, expression, or heme insertion. Together, these data demonstrate *S. oneidensis* can be optimized for OmcZ production and provide mechanistic insight into the conditions best suited for expression of OmcZʹ.

### Engineered OmcZʹ for improved electrical performance

Next, we examined whether we could genetically control the redox and conductive properties of nanowires by introducing substitutions near the heme binding sites of OmcZʹ. Previous studies with peroxidases^23,24,25^ and myoglobin^26,27^ have shown that carboxylate-bearing residues (aspartate, glutamate) that form hydrogen bonds with an iron-coordinating histidine lead to its partial deprotonation and stabilization of Fe(III), thereby significantly lowering heme redox potential. Using the cryo-EM structures of OmcZ, PDB:8D9M^28^ and PDB:7LQ5^3^ as guides, residues proximal to the N_δ1_ of bis-histidines were selectively mutated to either Asp or Glu. A summary of selected mutations (**Fig.S10**) and their positions along the WT OmcZ heme array is shown in **Fig.2f**. Mutations were introduced into the StrepII OmcZʹ operon, **Fig.1e**, through overlapping primer pairs (**Table S3**) and expression confirmed by OmcZʹ protein capture from supernatants. Of the substitutions attempted, extracellular expression of OmcZʹ was observed in 14 of 21 initial point mutations. Successful substitutions were primarily observed when starting with polar, uncharged residues; when hydrophobic residues were substituted there was a general decrease in protein expression, possibly due to destabilization and degradation of OmcZʹ.

To determine whether mutations impacted the electronic performance of the nanowires, whole cell conductance assays with OmcZʹ variants were carried out on 200µm interdigitated electrodes (**Fig.2g**). OmcZʹ nanowires were expressed from the full operon (**Fig.1b**) in the *Shewanella* Δ4 strain^29^, a strain previously engineered by our group to lack native filaments (including typeIV pili, mannose-sensitive pili, and flagellum). The current response from potential sweeps (IV curve) of the *Shewanella* Δ4 base strain revealed low conductance compared to strains expressing OmcZʹ and OmcZʹ variants, indicating expression of OmcZʹ led to a significant improvement in whole cell electronic performance. The individual substitutions I13E, T82D, V120D, and T173D showed conductance improvements over wild-type OmcZʹ (**Fig.2h**), with T82D having a roughly 1.5-fold increase in whole cell conductivity. Successful individual variants were then combined into double, triple, and quadruple mutants, with whole cell conductance measurements (**Fig.2g-h**) highlighting the improved conductivity of select double (OmcZʹ[I13E,T173D] and OmcZʹ[I13E,V120D]), triple (OmcZʹ[I13E,V120D,T173D], and quadruple (OmcZʹ[I13E,T82D,V120D,T173D]) mutants. Of the OmcZʹ variants tested, the quadruple mutant OmcZʹ[I13E,T82D,V120D,T173D] displayed the highest conductivity, with a nearly 2-fold increase relative to wild-type OmcZʹ. Improvements in conductance were roughly stepwise, suggesting that the variants are acting additively.

### Phylogenetic mining identifies novel structural features for OmcZʹ engineering

Having shown that whole cell conductance could be impacted by sequence substitutions, we carried out a phylogenetic search for potentially improved OmcZ variants. Starting from OmcZ sequences biomined from phylogenetic data^3^, of 146 reference clusters from NCBI with 90% cluster identity to *G. sulfurreducens* OmcZ (**Fig.S11**), only 15 were found to utilize an atypical heme II binding loop. Specifically, we identified a homologue of OmcZ from *Methanogaster_A* sp. (**Fig.2i**) that utilizes a typical motif at heme II, positioning a tyrosine in contact of both hemes II and III in place of the native Pro-Pro turn that precedes atypical heme II insertion in *G. sulfurreducens* WT OmcZ. Based on this predicted structure, we hypothesized that substitution of residues 42-57 (Ala-Leu-Ile-Asn-Thr-Val-Thr-Pro-Pro-Ala-Ser-Cys-Ile-Asn-Cys-His) in OmcZʹ with the *Methanogaster_A* sequence 41-53 Met-Leu-Val-Gln-Thr-Ser-Tyr-Ser-Cys-Thr-Asp-Cys-His motif might promote a unique tyrosine-heme electronic interaction while also stabilizing this region. Accordingly, OmcZʹ residues A42M, N45Q, V47S, T48Y, and N55D were mutated while residues 49-51 (Pro-Pro-Ala) were deleted. Residues I44 and I54 in OmcZʹ, corresponding to V43 and T50 in the homolog, respectively, were left unchanged due to their role in inter-subunit association. Thus, we constructed a chimeric variant of OmcZʹ, OmcZʹ[I13E,A42M,N45Q,V47S,T48Y,Δ49-51,N55D,T82D,S119D,V120T], hereafter referred to as OmcZ^+^ (**Fig.2i, Fig.S12**).

To better assess the impact of this novel domain on electrical performance we measured the bulk conductivity of 5µg purified OmcZʹ nanowires assembled from purified components on 5µm interdigitated electrodes (**Fig.S13**). Purified nanowires from OmcZ^+^ displayed the highest conductance of all tested substitutions (**Fig.2j**) with a roughly 3.5-fold increase over WT OmcZʹ (**Fig.2k**). Underlying double and triple variant conductance increased additively, as previously observed in whole cell measurements (**Fig.2h**). With actual OmcZʹ yields decreasing with added substitutions (**Fig.S13**), expression changes between the variants likely account for the differences in conductivity enhancements observed between whole cells and *in vitro* assembled and purified nanowires.

To determine whether enhanced conductivity was accompanied by changes in the intrinsic redox properties of the nanowires, we next analyzed isolated OmcZʹ variants via cyclic voltammetry. Isolated OmcZʹ nanowires were resuspended anaerobically to 40µM in 0.1X PBS pH7.4 (to impede sample aggregation^3^) and electrochemical performance was measured on gold screen-printed electrodes (SPEs) with an Ag reference (**Fig.2l**). Importantly, peak positions remained largely unchanged across all variants, suggesting OmcZʹ tertiary structure remains largely intact throughout the tested OmcZʹ variants, an observation supported by TEM imaging (**Fig.S14**) of purified, formed nanowires. That said, peak current amplitudes increased for engineered OmcZʹ variants. The mutation T82D, which is adjacent to solvent inaccessible hemes I & IV, may actively influence internal heme-heme electron redistribution, as evidenced by a peak shift from -140mV in samples lacking the mutation to -65mV in OmcZʹ[I13E,T82D,V120D,T173D] and -76mV in OmcZ+ (**Fig.S15**). In addition, in OmcZ^+^ -- the only variant containing a tyrosine residue proximal to the heme cofactor -- an additional redox feature emerges near +140 mV that may reflect perturbation of the local heme redox environment or participation of a tyrosine-associated redox process. The observation in all samples of a dominant anodic peak near -85mV following full reduction suggests that oxidation of OmcZʹ (and it’s variants) occurs cooperatively within a narrow potential window, potentially through solvent-accessible hemes that serve as electronic exit sites for electrons redistributed from the internal heme chain. This observed redox asymmetry is in contrast to other multiheme cytochromes such as MtrC and OmcA that display reversible and highly symmetric voltametric signals^30^.

### Cryo-EM structure of purified OmcZʹ and OmcZ^+^ nanowires

To evaluate whether heterologous expression in *S. oneidensis* preserves the structural features associated with the exceptional conductivity of OmcZ nanowires, cryo-EM analyses were carried out with purified OmcZʹ and OmcZ^+^ nanowires. Two-dimensional class averages of both OmcZʹ and OmcZ^+^ nanowires revealed filament morphologies highly consistent with those reported for native OmcZ (PDB: 7LQ5), including the characteristic sinusoidal nanowire architecture and periodic subunit organization (**Fig.3a**). Structures were fit to 2.9 Å resolution for OmcZʹ (**Extended Data Fig.1**) and 3.4 Å resolution for OmcZ^+^ (**Extended Data Fig.2**) nanowires following helical refinement in cryoSPARC and structural refinement against density modified half maps generated in Phenix. At these resolutions backbone densities, bulky sidechains, and all 8 hemes (including positions of extending propionate groups) could be confidently traced. Structural alignment of the heterologously expressed OmcZʹ with WT OmcZ (**Fig.S16**) revealed strong conservation of protein folding, including residues forming the heme II attachment site following native atypical motif mutation to the CXXCH motif (**Fig.3b**). Importantly, our removal of these residues did not significantly impact the OmcZ tertiary fold or heme arrangement. Nonetheless, closer examination of the cryo-EM density maps did identify some localized conformational differences between OmcZʹ and WT OmcZ (PDB:7LQ5). In particular, multiple proline residues within OmcZʹ exhibited cis conformations (20%, **Fig.3c**), corroborating the role of the prolyl isomerase in the proper folding and expression of OmcZ’ (**Fig.2e**).

**Figure 1.**
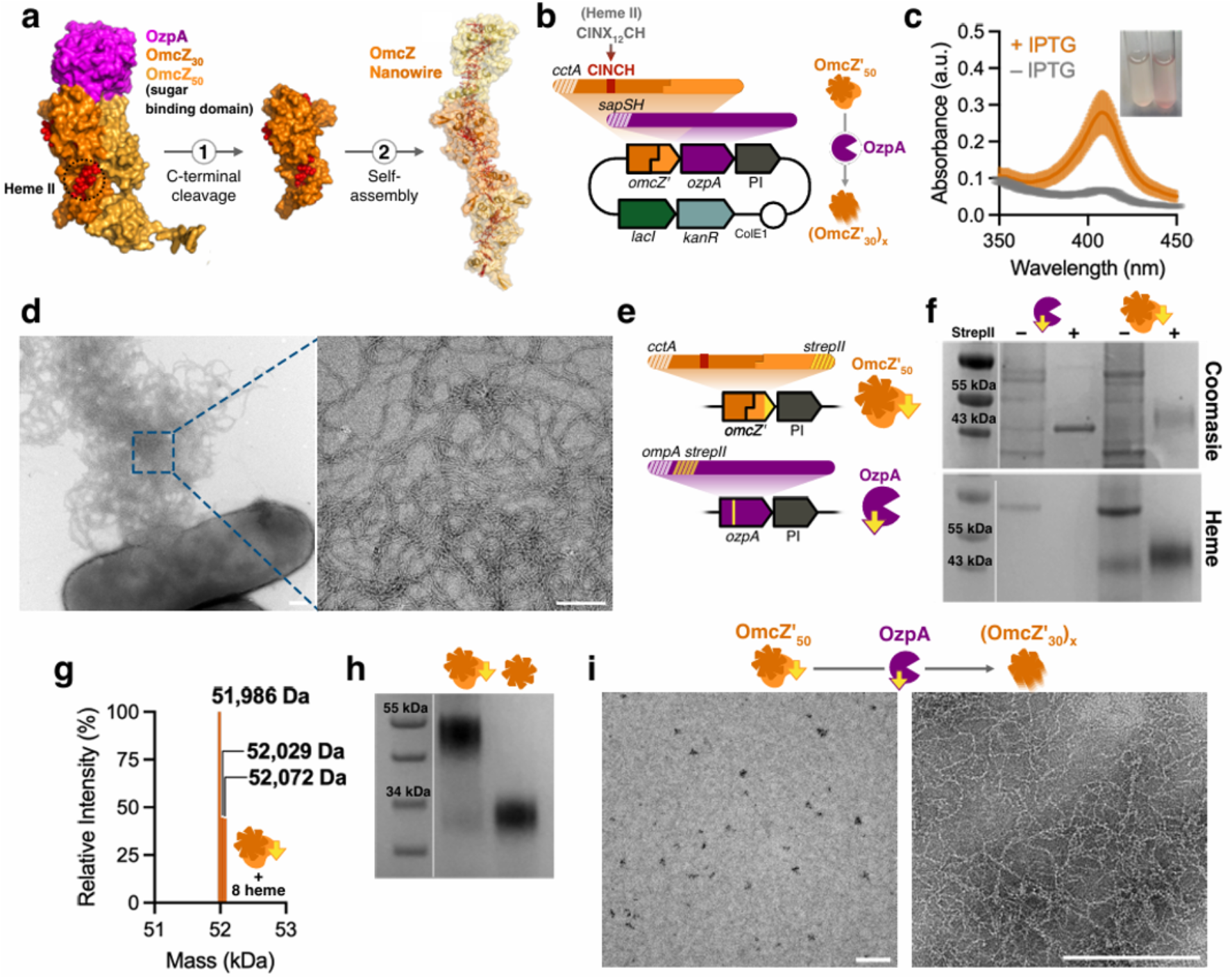
Expression of the octaheme cytochrome OmcZʹ in *Shewanella oneidensis.* (a) Schematic of OmcZʹ_50_ cleavage by OzpA in nanowire formation. (b) Plasmid map of 3 gene operon for *S. oneidensis* OmcZʹ *in vivo* nanowire expression. (c) Supernatant absorbance of induced (orange) and uninduced (grey) OmcZʹ expression; inset, Hungate tube *S. oneidensis* cultures following expression. (d) TEM of extracellular OmcZʹ nanowire formation in *S. oneidensis* MR-1 expressing OmcZʹ operon. (e) 2 gene operon design for affinity purification and *in vitro* nanowire formation. (f) Coomassie and heme stain of OzpA (lanes 1,2) and OmcZʹ (lanes 3,4) pre/post purification. (g) Weight determination by MS of strepII OmcZʹ. (h) Heme stain of purified OmcZʹ before (lane 1) and after (lane 2) incubation with OzpA. (i) TEM of OmcZʹ_50_ pre (left) and post (right) incubation with OzpA. All scale bars 200nm.

**Figure 2.**
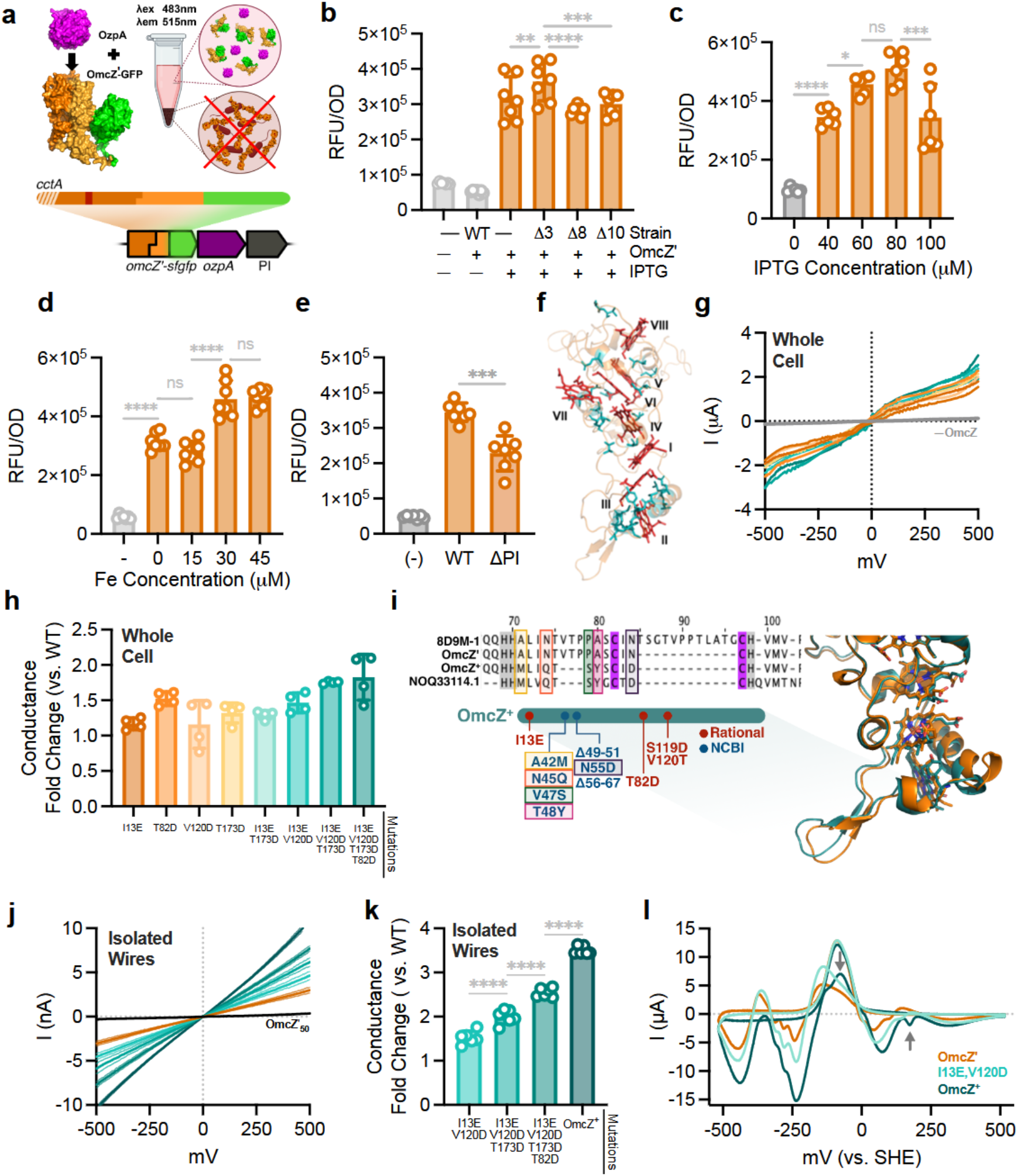
OmcZʹ optimization. (a) OmcZʹ-GFP fusion fluorescence from culture supernatant. (b) OmcZʹ-GFP supernatant fluorescence in *S. oneidensis* strains MR-1, JG596 (Δ3), JG1194 (Δ8), and JG1519 (Δ10). (c) Supernatant fluorescence of OmcZʹ-GFP in strain JG596 under varying IPTG induction. (d) Supernatant fluorescence of OmcZʹ-GFP in strain JG596 under varying iron supplementation. (e) Supernatant fluorescence of OmcZʹ-GFP in presence vs absence of prolyl isomerase. (f) Native OmcZ (PDB:8D9M, orange) with heme array (red) and rational mutations (teal). (g) Whole cell IV curves of *Shewanella* Δ4 strains expressing OmcZʹ variants on 200µm IDEs. (h) Bar graph of conductance fold increase in engineered OmcZʹ variants vs WT OmcZʹ. (i) OmcZ^+^ homology (left) and predicted structure (teal,right) overlayed on WT OmcZʹ (orange). (j) IV curves of *in vitro* assembled OmcZʹ nanowires on 5µm IDEs. (k) Bar graph of conductance fold increase in engineered OmcZʹ variants vs WT OmcZʹ. (i) Cyclic voltammetry of purified WT and engineered OmcZʹ nanowires on gold screen printed electrodes. Arrows denote T82D peak shift, top, and Tyr-heme peak, bottom, engineered into OmcZ^+^. (b-e, k) Significance testing performed using one-way ANOVA, *p<0.05, **p<0.01, ***p<0.001, ****p<0.0001

Despite these local differences, the overall filament architecture remained highly conserved. OmcZʹ exhibited helical rise and twist parameters nearly identical to those of WT OmcZ (Δz = 57.6 Å, Δφ = −158.9° versus Δz = 57.4 Å, Δφ = −159.1° for 7LQ5), whereas OmcZ^+^ retained a similar rise but displayed a more pronounced change in helical twist (Δz = 57.8 Å, Δφ = −170.0°; **Fig.3d**). Edge-to-edge (ETE) heme distances, and Fe-Fe spacing within the linear heme stack varied only modestly among the three structures, consistent with preservation of the overall filament architecture (**Fig.3e**). Together these results demonstrate that heterologous expression in *S. oneidensis* and mutagenesis of key regions do not significantly disrupt nanowire structure and preserve contiguous electron hopping between hemes.

### OmcZ^+^ nanowire architecture and inter-subunit heme interactions

While the overall structure of OmcZ^+^ was similar to the wild-type (**Fig.3d–e**), some local structural reorganizations were notable. Heme pocket II–VIII (S119D,V120T; **Fig.S17**), IV (I13E; **Fig.S18**), and I–IV for T82D (**Fig.3f**) had increased hydrogen bonding interactions and decreased carboxylic proximity to the N_δ_1 of heme-coordinating histidines. Tyr48 in particular proved to be in close-atom contact with Hemes II-III (**Fig.3g**), in close agreement with the predicted models (**Fig. 2i**).

More substantial inter-subunit differences resulted from replacement of the Heme II binding region (ALINTVTPPASCINCH) with the homolog-derived sequence (MLIQTSYSCIDCH) in OmcZ^+^. OmcZ^+^ displayed subunit interfacing involving heme binding domains VII-II as well as the traditional VIII-II heme binding domain interactions that define linear nanowire assembly, resulting in the formation of “dendrite-like” or branched nanowires. Separately, pentameric ring structures (**Fig.3h**) were also observed. These structures had a preferred “face” orientation bias during freezing, making high resolution map reconstruction near impossible without high volume tilt data. Accordingly, we constructed a lower-resolution map using C5 symmetry for rough fitting of OmcZ^+^ density. Reconstruction revealed assembly of radially arranged OmcZ^+^ subunits with consecutive Heme VII-II interfaces, forming a closed ring with an open central cavity. Additionally, members of the pentameric ring could be fit with a second OmcZ^+^ subunit at the traditional Heme VIII-II interface (**Fig.3i**), forming nanowire extensions at each of the 5 pentamer subunits, with the lengths of these extensions and further branching varying. Notably, OmcZ^+^ displayed several close atomic interfaces involving Hemes VII–II (**Fig.3j**), while fitting of OmcZ’ with pentamer map density led to strong inter-subunit clashes. The relative abundance of linear (Heme VIII:II) and dendritic (Heme VII:II) morphologies were determined by direct inspection of OmcZ^+^ micrographs (**Fig.S19**). Linear nanowire assembly remained the dominant morphology (∼58%), while dendritic assembly accounted for ∼42% of observed subunit interfaces (**Fig.3k**). Pentameric rings were also frequently observed, occurring at an average of ∼16 assemblies per micrograph (range 8–23, **Extended Data Fig.3**).

**Figure 3.**
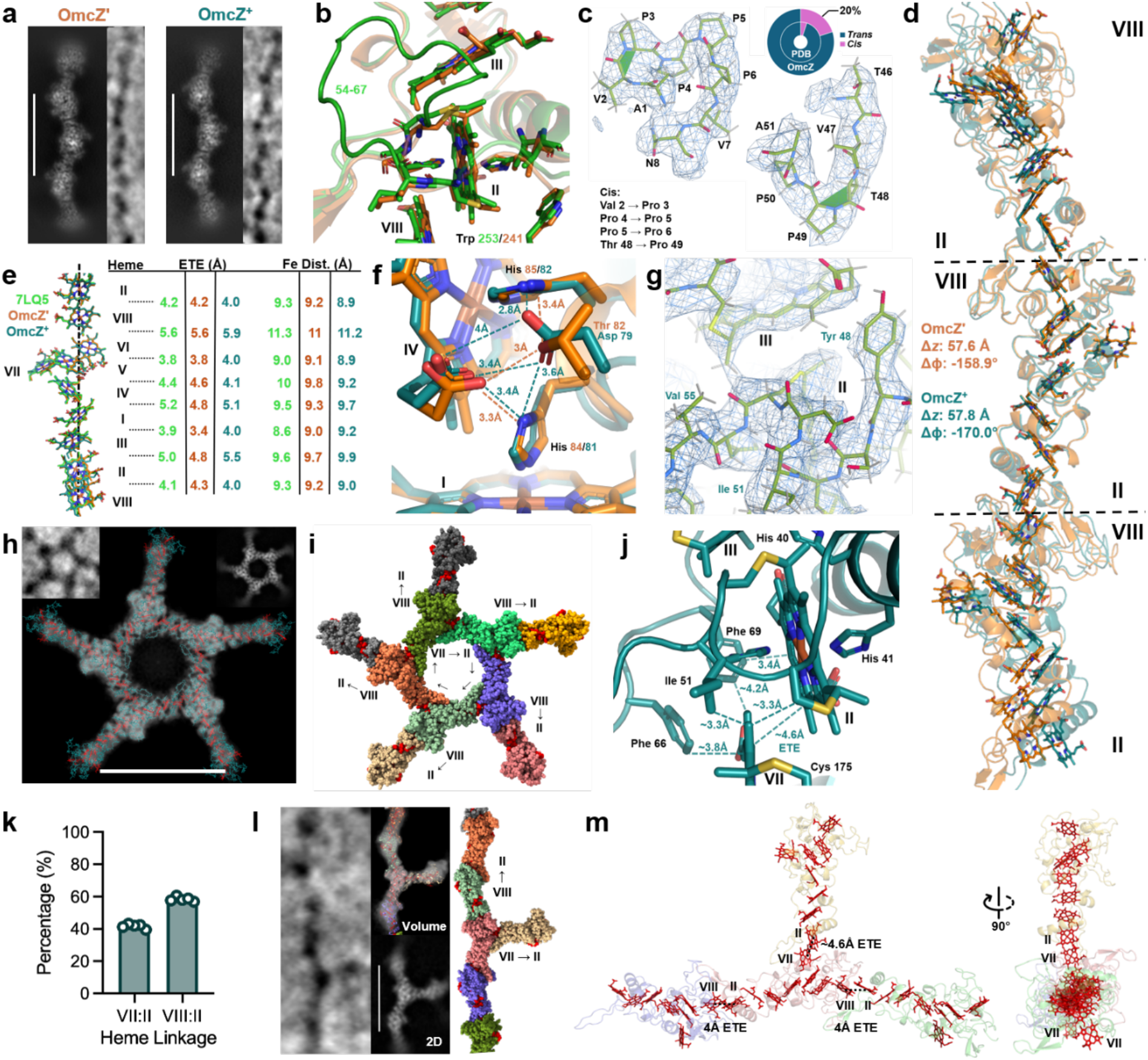
OmcZ^+^ structure influences unique heme architecture. (a) 2D class averages of OmcZʹ and OmcZ^+^ nanowires. (b) Alignment of OmcZ (*G. sulfureducens*, PDB 7LQ5, green) with OmcZʹ (*S. oneidensis*, orange). (c) Cryo-EM density fitting of cis conformation prolines (20%, inset) in OmcZʹ. (d) Alignment of OmcZʹ (orange) and OmcZ+ (teal) nanowires. (e) Edge-to-edge and iron center distances of OmcZ nanowire (OmcZ:7LQ5, OmcZʹ:36HZ, OmcZ^+^:36KE) linear heme chain. (f) T82D alignment and hydrogen bonding of Heme I-IV pocket of OmcZʹ (orange) and OmcZ^+^ (teal). (g) Cryo-EM density illustration of OmcZ homolog mutation in OmcZ^+^. (h) OmcZ^+^ pentamer volume map with OmcZ^+^ nanowire dimers (36KE; chain, teal; heme, red) fit to map density. Inset, micrograph raw particle (top left) and sample 2D class average (top right). (i) OmcZ^+^ pentamer surface with individual subunits (colored) and hemes (red). (j) OmcZ^+^ Heme VII-II subunit arrangement with close atom contacts. (k) Graph of OmcZ^+^ nanowire assembly as observed in rough inspection of micrographs. (l) Dendritic assembly in OmcZ^+^. Representative micrograph raw particle (left), sample 2D class average (bottom middle), structural interpretation fit to map density (top middle), surface map of structural interpretation (right). (m) Internal heme chain (red) from fit model (l). Scale bars 120Å.

To further investigate the structural basis of branching, ∼20,000 particles were used to reconstruct a low-resolution map of a representative OmcZ^+^ dendritic nanowire, which was subsequently fit with a model derived from OmcZ^+^ (36KE) and the Heme VII-II dimer interface (**Fig.3l-m**). The fitted dendritic reconstruction was consistent with the structural interpretation derived from the structures of OmcZ^+^ nanowires and Heme VII-II dimers, ultimately supporting a model in which alternative Heme VII-II interfaces generate branched filament architectures observed directly in cryo-EM micrographs (**Extended Data Fig.4**). Ultimately, altered interfacial interactions of OmcZ^+^, together with the enhanced conductivity observed from several single-point mutations, suggest that subtle changes in heme environment and inter-subunit organization may influence both intra- and intermolecular charge transport within OmcZ nanowires.

**Figure 4.**
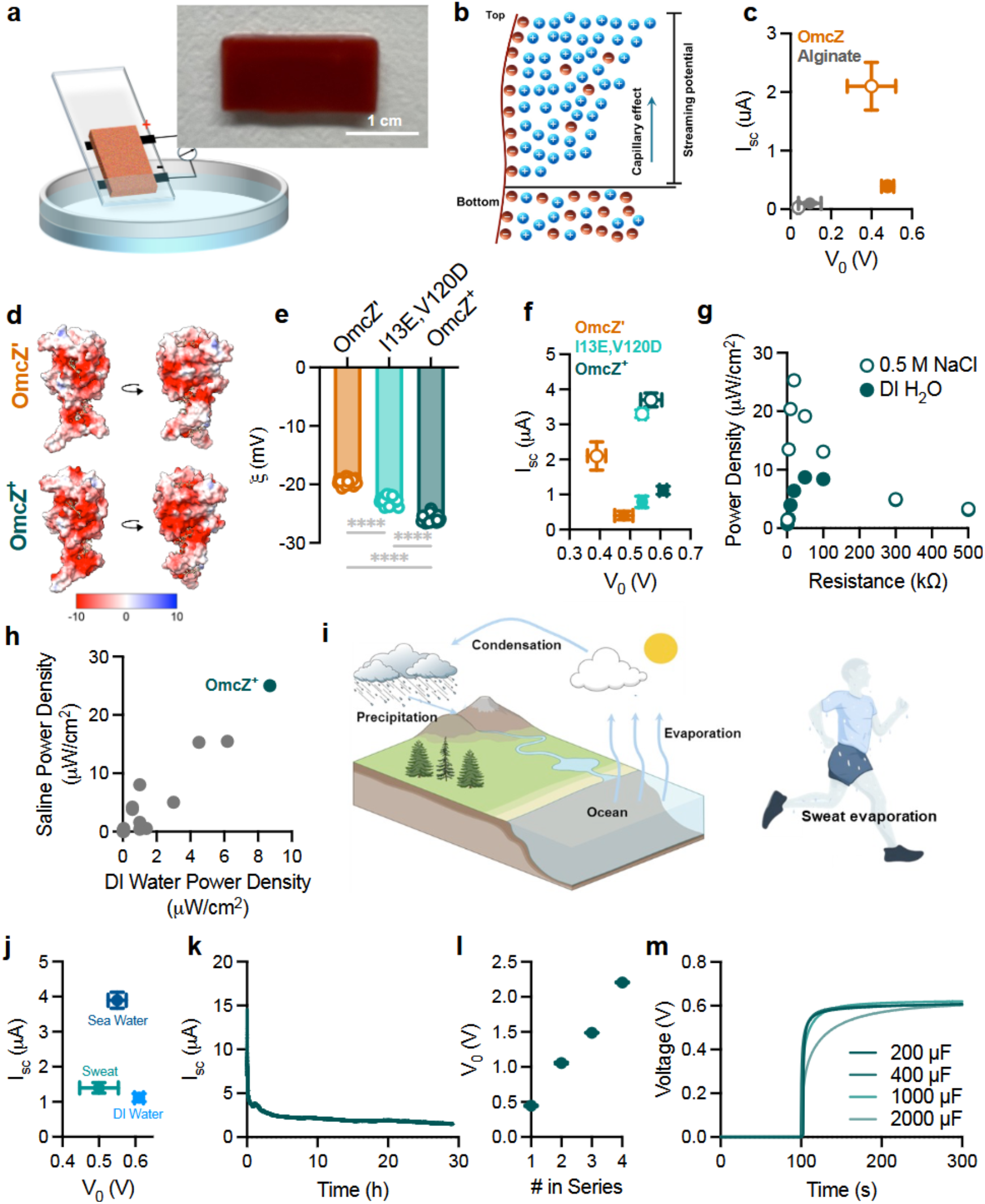
Hybrid OmcZʹ::alginate hydrogels for hydrovoltaic application. (a) Top, Photograph of the as-fabricated OmcZʹ::alginate hydrogel (right panel). Bottom, diagram of the device structure. (b) Schematic of evaporation-enabled electricity generation in OmcZʹ::alginate hydrogel. (c) Hydrogel voltage and current performance in deionized (DI, solid points) and salt (0.5M, hollow points) water. (d) Electrostatic surface map of WT OmcZʹ (top) and OmcZ+ (bottom). (e) Zeta potential of formed nanowires (OmcZ’_30_) of OmcZʹ and mutants. (f) Performance of OmcZʹ hydrogel devices in DI water (solid) and 0.5 M NaCl (hollow). (g) The relation between power density and load resistance of OmcZ+ hydrogel device in both DI water and 0.5 M NaCl conditions. (h) The performance comparison chart of OmcZ+ hydrogels in relation to other evaporation-enabled electric generators^34,38,41,44,45,46,47,48,49,50,51,52,53^. See Table S4 (i) Water evaporation diagram. (j) Current and voltage outputs of the OmcZ+ hydrogel device in different solutions. (k) A continuous recording of the short-circuit current from OmcZ+ hydrogel for >30hrs. (l) Voltage performance increase with OmcZ+ hydrogel devices in series. (m) Charging curves of a single OmcZ+ hydrogel device in 0.5 M NaCl solution for capacitors with different capacitances.

### Evaporation induced energy generation using OmcZʹ

Ambient water represents a ubiquitous, continuously available, and sustainable energy resource^31,32,33^ and protein nanowires from *G. sulfurreducens* have emerged as promising materials for such devices due to their exceptional electronic properties^34,35,36^. Despite these advantages, previous efforts at incorporating protein nanowires into evaporation-based energy generators have been limited by an inability to produce these materials at a reasonable scale or level of purity^37^. Enabled by our heterologous expression strategy, we explored the use of OmcZʹ nanowires, and their variants, in water evaporation-induced electricity generators (WEG). To enhance mechanical robustness while maintaining flexibility for future integration with wearable devices (**Fig.S20**), we embedded OmcZʹ nanowires (2 wt%) in an alginate biopolymer matrix, with the macroscopic geometry defined by a 3D-printed mold (**Fig.S21**). We first evaluated electricity generation from water evaporation using a device fabricated by placing two carbon-cloth electrodes across a prepared hydrogel film supported on a glass substrate (**Fig.4a-b**). Upon immersing one terminal in DI water, OmcZʹ-containing devices generated a spontaneous voltage of ∼0.48 V and sustained a steady current of ∼400 nA (**Fig.4c**). Alginate-only devices generated minimal voltage and current. The maximum power density generated (∼4.7 μW/cm^2^) following OmcZʹ incorporation was nearly two orders of magnitude larger than that previously achieved with carbon and other biologically derived materials^38,39^.

Power generation in evaporation devices is driven by anisotropic ion transport in the interfacial electric double layer, giving rise to a streaming potential across the electrodes (**Fig.4b**). Accordingly, current generation is directly proportional to the surface charge of ion transport channels within the hydrogel matrix^40^. We predicted that both the I13E,V120D OmcZʹ mutant and OmcZ^+^ would exhibit more negative surface potentials and superior device performance. Indeed, zeta potential measurements of OmcZʹ_30_ wires for I13E,V120D and OmcZ^+^ showed significantly more negative zeta potentials relative to WT OmcZʹ across multiple pH conditions (**Fig.4e, Fig.S22**). As predicted, genetically encoded changes to surface potential were accompanied by improved evaporation-enabled power generation. Specifically, devices incorporating I13E, V120D and OmcZ^+^ wires sustained steady currents of 0.8 and 1.1 uA, respectively (**Fig.4f**), almost double the performance of WT OmcZʹ wires.

Next, we examined the effect of salt concentration on current generation. High salt concentrations typically suppress the electrical output of evaporation-driven generators (often to <5% of that in pure water), because the increased ionic strength shortens the Debye length, weakens electric double-layer overlap, and thereby lowers streaming efficiency^41^. In contrast, all hydrogels containing OmcZʹ showed improved performance in saline media (0.5 M NaCl). Although generated voltage decreased slightly, output currents increased to 2 uA, 3.5 uA, and 4 uA for WT OmcZʹ, the I13EV120D mutant, and OmcZ^+^ respectively. Notably, the power–resistance characteristics in OmcZ^+^ hydrogels were further enhanced in saline media with reduced internal resistance (∼20 kΩ) and increased maximum power (∼25.3 µW/cm^2^), approximately threefold higher than the same device in DI water (**Fig.4g**). Benchmarking against previously reported evaporation-enabled generators, including those fabricated using whole *Geobacter* biofilms, places OmcZ^+^ devices among the top-performing systems, combining competitive power density with stable operation in high-salinity environments (**Fig.4h, Table S4**).

Finally, we evaluated the performance of devices containing OmcZ^+^ in representative real-world aqueous environments (**Fig.4i)** and assessed their scalability for practical energy storage and utilization. Devices containing OmcZ^+^ nanowire hydrogels delivered stable electrical outputs using multiple solutions and produced an open-circuit voltage of ∼0.51 V and a short-circuit current of ∼1.4 μA in human sweat (ISO 11641), and ∼0.55 V and ∼3.9 μA in seawater (ASTM D1141-98), respectively (**Fig.4j**). In sea water, devices maintained a steady current output (∼1.5 μA) for over 30 hours (**Fig.4k**). Power output could be modularly scaled through device integration and four devices in series increased the total voltage approximately linearly to ∼2.4 V (**Fig.4l**). In addition, a single OmcZ^+^ protein nanowire hydrogel device was able to charge commercial capacitors (**Fig.4m**), with all capacitors reaching a stable voltage of ∼0.55 V. Notably, a 2000 μF capacitor charged in parallel reached 0.55 V within 2 min, and the stored energy could directly power an LED. Together, these results highlight the practical promise of OmcZʹ hydrogels as salt-tolerant, scalable power sources for self-powered electronics in diverse aqueous environments.

## DISCUSSION

Our ability to express *Geobacter* OmcZ in a more tractable *S. oneidensis* host overcomes several challenges that have previously hindered the characterization and application of cytochrome nanowires. In contrast to *E. coli*, *S. oneidensis* natively produces several cytochromes containing 4 or more hemes and is well-suited to the heterologous expression of multiheme proteins. When transformed with a plasmid containing the full OmcZ operon (OmcZ, OzpA, prolyl isomerase), *S. oneidensis* strains formed OmcZʹ nanowires under typical anaerobic growth conditions. We found that the prolyl isomerase was required for OmcZ nanowire expression while a fourth gene within the operon for other Omc nanowires, which encodes a highly repetitive protein structure, was not required for nanowire formation^42^. While OmcZʹ expression is robust in *S. oneidensis*, we observed no effect on EET flux, suggesting that the nanowires are not connected to *S. oneidensis* redox metabolism.

Since nanowire purification from *G. sulffureducens* cultures is notoriously difficult and samples are often contaminated with pili, other cytochromes, and various cellular components, an *in vitro* assembly method from purified proteins is practically required for future biochemical and biophysical studies. The native operon was split into two different *S. oneidensis* strains, one expressing OzpA and the other OmcZʹ. Following separate purification of OzpA and OmcZʹ_50_ from these strains the two proteins could be added together *in vitro* to form highly pure OmcZʹ_30_ nanowires. This expression system produced highly pure nanowire formulations and should aid in more comprehensive studies of OmcZʹ polymerization and nanowire dynamics.

Genetic control over nanowire formation both in cells and *in vitro* ultimately allows facile strain engineering and rational mutagenesis of OmcZʹ. When residues predicted to influence heme redox potentials were varied, 14 of 21 single substitutions resulted in successful OmcZʹ expression, with several single substitutions producing measurable improvements in bulk conductance with both whole cell and purified nanowires. More importantly, by using the wealth of phylogenetic data available from genomic and metagenomic databases, we identified naturally occurring structural motifs from divergent OmcZ homologs. Further transplanting one such element, the heme II loop from *Methanogaster_A* sp., into the *Geobacter sulfurreducens* OmcZ scaffold, led to OmcZ^+^, which exhibited bulk conductivity approximately 3.5-fold greater than that of wild-type OmcZʹ. The successful incorporation and assembly of this foreign structural element, coupled with the retention of function across numerous engineered conductance mutations, demonstrates an unexpected degree of modularity within OmcZ nanowires and suggests their architecture can tolerate substantially more sequence diversity than previously understood. This result establishes a framework for the further design of chimeric cytochrome nanowires with other functional fusions.

Given the sequence divergence of OmcZ^+^ from its parents, we evaluated the degree to which resultant wires were structurally similar to or distinct from those derived from the parental OmcZʹ. The Fe-Fe distance, edge-to-edge distance, wire helicity, and other structural parameters were largely similar between OmcZʹ and OmcZ^+^. Our solved structures also nicely explain the role of the prolyl isomerase in OmcZ expression: in contrast to the previously solved structures of OmcZ nanowires^3,28^, our structure contains 20% of cis-prolines, a value significantly higher than the average in the PDB (∼5%). Several purported OmcZ operons do not encode a prolyl isomerase, suggesting the testing of additional variants could further illuminate wire assembly.

In contrast to OmcS and OmcE, OmcZ possesses one solvent-exposed or “leaky” heme (Heme VII). The structures of OmcZ^+^ provide insight into the potential role of this heme in nanowire construction and assembly. We observed unique “dendrite-like” and pentameric ring structures during OmcZ^+^ wire formation. In both dendritic and ring structures, Heme II and Heme VII form the key interface between adjacent subunits. While OmcZʹ also sometimes adopted the Heme VII-II dimer interface, **Fig.S23**, its prevalence was far lower (<5%) than that for OmcZ^+^ (∼42%). Whether these structures are representative of those that can be formed by native OmcZ or its homologs, or are synthetic artifacts from our mutational studies and *in vitro* assembly conditions, will require further investigation. Regardless, our results along with recent evidence for bundled OmcE wires in the literature^43^, suggest that cytochrome nanowires may adopt a wide range of non-linear morphologies. Further characterizing the electromagnetic properties of these novel structures will require careful single wire measurements, which should be facilitated by our ability to produce pure wire formulations.

*Geobacter* pili have previously been used as evaporation-induced electricity generators^35^. Water evaporation-induced electricity generators incorporating OmcZʹ nanowires generated a maximum power density of ∼4.7 μW/cm² in pure water, nearly two orders of magnitude higher than previously reported devices of similar design. Notably, in contrast to many evaporation-driven generators, OmcZʹ protein nanowire hydrogels maintained robust electrical output in saline environments. Incorporation of engineered variants OmcZʹ[I13E,V120D] and OmcZ^+^ further enhanced both voltage and current outputs, likely reflecting changes in charge density and ion transport arising from modifications to the protein structure. In particular, OmcZ^+^ increased the maximum power density to ∼25.3 μW/cm². We also examined device performance under representative real-world conditions and OmcZ^+^ nanowire devices were tested in seawater and human sweat, producing short-circuit currents of ∼3.9 μA and ∼1.4 μA, respectively. Device performance in seawater remained stable for days, and multiple devices could be connected in series to increase voltage output approximately linearly. The energy generated by a single device was sufficient to charge a 2000 µF capacitor in under two minutes, enabling direct powering of an LED. Together, these results highlight the potential of protein nanowire hydrogels as a salt-tolerant and scalable power source for self-powered electronics operating in diverse aqueous environments.

Overall, our expression system can be leveraged to rapidly examine OmcZ homologs and novel operon structures, while the ability to form nanowires *in vitro* should enable more robust biochemical and biophysical characterization of these unique protein materials. Further improvements in expression and assembly should allow scaling for routine device fabrication. Most importantly, our results demonstrate that OmcZ, and it’s homologs, can form unexpected supramolecular structures. Together with our ability to genetically engineer conductivity, these structures provide a tantalizing glimpse into the potential construction of genetically-defined molecular electronics.

## MATERIALS AND METHODS

### *S. oneidensis* bacterial strains and expression vectors

*Shewanella oneidensis* MR-1, Δ4^29^, JG596^15^, JG1194^16^, and JG1519^16^ strains were all transformed with plasmid backbones originating from pESOpWT^29^ containing ColE1 origin of replication, kanamycin resistance, and IPTG inducible LacI promoter/repressor. Native *omcZ*, *ozpA*, and prolyl isomerase genes were ordered as *E.coli* optimized gBlocks (IDT) with N-terminal signal sequences and BsaI overhangs (**Table S2**). pESOpWT backbone was amplified using primer pair listed in **Table S3**. The 3 genes and pET backbone were assembled by golden gate and transformed into electrocompetent *S.oneidensis* MR-1 by electroporation at 1.25keV in BIO-RAD pulsar. The resultant native OmcZ plasmid, pESoZgs , was further modified by primer pair (**Table S3**) to mutate native heme II loop (CINTSGTVPPTLANKGCH) to a typical heme motif (CINCH). The amplified linear fragment was assembled by golden gate and transformed into electrocompetent *S. oneidensis* MR-1 as mentioned above to produce plasmid pESoZʹ, **Fig.1b**, for OmcZ[Δ56-67] expression.

OmcZʹ-GFP expression plasmid was assembled using primer pairs listed in **Table S3** and utilized PCR amplified sfGFP gene from pKAR2^21^. Fragments and pESoZʹ vector backbone were assembled using golden gate and transformed into electrocompetent *S. oneidensis* JG596 and JG1519.

Plasmids for *in vivo* biofilm conductance, **Fig.1b**, and *in vitro* strepII affinity chromatography, **Fig.1f**, were obtained by PCR amplification of primer pairs listed in **Table S3,** assembly using golden gate, and electroporation into electrocompetent *S. oneidensis* strain Δ4 or JG596, respectively.

Individual OmcZʹ mutations were made from primer pairs listed in **Table S3** and amplified within OmcZʹ strepII expression vector, **Fig.1b**. For double, triple, quadruple, and OmcZ^+^ mutations, primer pairs were used on the corresponding plasmid backbones from previous mutations.

### *S. oneidensis* growth and expression conditions

*S. oneidensis* strains were streaked on LB plates containing 25 µg/mL Kanamycin antibiotic as appropriate from glycerol stocks (18% glycerol) and incubated overnight at 30C. For OmcZʹ and OzpA expression, individual colonies were picked from overnight streak plates and grown in LB broth containing 60mM HEPES pH 7.4 and 0mM salt to an OD_600_ of 1.5-2 in 50mL sealed Eppendorf tubes in 30C shaking incubator at 220RPM. Anaerobic expression media was prepared using Shewanella Basal Media (SBM) modified with 0.5g/L yeast extract, 3g/L casamino acids, 60mM HEPES pH7.6, 60mM Sodium Fumarate, 1mL/L ATCC vitamin solution, 1mL/L ATCC trace minerals, 30µM FeCl_3_ (OmcZʹ expression only), 20µg/mL Kanamycin (as appropriate), and 100µM IPTG (as appropriate) in membrane sealed bottles. Prepared media was made anaerobic by purging under 10PSI argon flow for 20min/L. Upon reaching OD_600_, cultures were centrifuged at 4,200 *x g* room temperature and supernatant discarded. Cell pellets were resuspended with anaerobic media, syringe injected into anaerobic expression media, and bottle membranes covered with mineral oil. Anaerobic cultures were placed in standing 30C incubators for 24 hours for protein expression.

For OmcZʹ-GFP expression, individual colonies were picked from overnight streak plates and grown in LB broth containing 60 mM HEPES pH 7.4 and 0 mM salt for 10 hours in 96-well deep well plates (0.5 mL) in 30C incubator shaking at 220 rpm. Anaerobic expression media was prepared using Shewanella Basal Media (SBM, ref) modified with 0.5g/L yeast extract, 3g/L casamino acids, 60mM HEPES pH7.6, 60mM Sodium Fumarate, 1mL/L ATCC vitamin solution, 1mL/L ATCC trace minerals, 30μM FeCl3 (OmcZ expression only), 20μg/mL Kanamycin (as appropriate), and 80μM IPTG (as appropriate) in membrane sealed bottles. Prepared media was made anaerobic by purging under 10PSI argon flow for 20min/L. After 10 hours, plates were spun down at 2200 *x g* for 20 minutes then all materials were transferred to the anaerobic chamber where cell pellets were resuspended with 1 mL anaerobic media (without IPTG) and allowed to set for 30-60 minutes. Cultures were then transferred at 1:100 volume to new plates containing fresh anaerobic media for induction (with IPTG). After 20 hours of growth, induction plates were removed from the anaerobic chamber and OD_600_ values were collected of individual wells. Plates were then spun down at 2200 *x g* for 20 minutes and 200 uL supernatant was transferred to 96-well plates followed by incubation at 4 °C overnight. The following day, RFU values were obtained of supernatant plates using 483nm excitation and 515nm emission.

### OmcZ_50_ and OzpA purification

Following 24 hour anaerobic expression cultures were centrifuged at 4,500 *x g* at 4C for 30min. Culture supernatant was run through 0.2 micron filters and 10ml/L protease inhibitor (EDTA free, Thermo Fisher) was added to supernatant of cultures expressing OmcZʹ. Additionally, OmcZʹ cell pellets were resuspended in 50mM HEPES pH8 at 100mL/L culture and stirred at 4C for 2 hours in a cold room to dissociate remaining OmcZʹ_50_ from cells. Following dissociation, resuspended cells were centrifuged at 8,000 *x g* at 4C for 30min. Resuspension supernatant was filtered through 0.2 micron membranes and added to previous culture supernatants.

StrepII affinity columns were prepared using high capacity strepII resin (31mg/mL binding capacity, IBA). Culture supernatants were run through appropriate columns and washed with 100 column volumes of wash buffer containing 50mM HEPES pH8, 120mM NaCl, 2% glycerol and 100µM EDTA. Elution was performed with appropriate wash buffer containing an additional 50mM biotin. Following elution, OmcZʹ samples were dialyzed twice at 4hrs minimum in 20mM HEPES pH7.2, 10mM NaCl, and 2% glycerol (1:100 volume minimum) at 4C and OzpA samples were dialyzed (20 kDa dialysis cassette) twice at 4hrs minimum in 20mM HEPES pH7.2, 20mM NaCl, and 2% glycerol (1:100 volume) at 4C. OzpA auto-maturation was carried out following dialysis by refreshing media and leaving overnight at room temperature. Following dialysis and maturation samples were collected and stored at 4C.

### *In vitro* OmcZ nanowire assembly

OmcZʹ samples were prepared to ∼5uM concentration in 20mM HEPES pH7.2, 10mM NaCl, and 2% glycerol and incubated with mature OzpA (∼500µg/mL) at 30:1 v/v. Samples were injected in 20 kDa dialysis cassettes and dialyzed in 20mM HEPES pH7.2,10mM NaCl, and 2% glycerol for ∼24hrs at room temperature. OzpA was removed from samples through first bringing solution to 50mM HEPES pH8 followed by addition of saturated ammonium sulfate for nanowire precipitation at 4C. OmcZʹ samples were then centrifuged at 10,000 *x g* for 10 minutes and supernatant removed. Nanowire pellets were resuspended in appropriate buffers for TEM imaging, electrical characterization, or cryoEM imaging, with additional rounds of dialysis performed as necessary to remove residual salts. OmcZʹ_50_ pro-peptide cleavage was confirmed by SDS PAGE gel staining and subsequent nanowire self-assembly through TEM.

### Negative stain TEM

400 mesh gold grids were plasma cleaned at 20mA for 30 seconds and 5uL samples were drop cast on the grids and absorbed for 2 min. Excess liquid was blotted by filter paper and grids were then washed with 20uL of pure H_2_O and blotted again. 5uL of 1% phosphotungstic acid (pH adjusted to 7) was then added to sample surface and stained for 30 seconds. Excessive stain was removed by filter paper and another 5uL of 1% phosphotungstic acid was added to the grids for 30 seconds followed by blotting. Grids were air dried before being transferred to TEM. TEM images were taken with JEOL 1400 S/TEM microscope at 120 kV operating voltage.

### Biofilm conductivity

*Shewanella oneidensis* Δ4 strain was transformed with OmcZʹ plasmids listed under **Table S2** containing the full OmcZʹ operon. Cells were grown and expressed, with antibiotic and IPTG as appropriate, under previously described conditions above in 250mL membrane sealed bottles. Following 24 hours of expression 50mL of sample was added to Eppendorf tube and centrifuged at room temperature at 4,200 *x g* for 10min. Supernatants were discarded and cell pellets were scraped with inoculation loops and placed onto plasma cleaned (20mA for 30 seconds) 200 µm gold interdigitated electrodes (Metrohm). Samples were allowed to air dry for 20 minutes before taking dual voltage sweeps from -650mV to +650mV on a Keithley 4200-SCS using 2 point probe setup under normal conditions at room temperature. 3 dual sweeps were performed starting from negative voltage and 3 dual sweeps performed beginning from positive voltage. Each individual run was then averaged and used to calculate overall conductance average for each sample.

### Phylogenetic tree

Sequences were sourced from the NCBI protein database using BLASTp against uniprot sequence 8D9M^28^. All sequences with a bitscore of >100 were selected. Outgroup sequences (OmcS family) were selected from an NCBI BLASTp search against NCBI accession WP_010943141.1. An alignment was then conducted using MAFFT and trimmed using TRIMAL. The alignment was visualized on Jalview and all OmcZ-like sequences missing the 8 heme binding motifs and distal HH pairs were removed from the alignment. A phylogenetic tree with curated sequences and outgroup sequences was run using IQtree with a bootstrap value of 1000. The phylogenetic tree was visualized on iTOL.

### Cyclic Voltammetry

OmcZʹ samples were prepared to 40µM concentration in 0.1X PBS at pH 7.4. Samples were then transferred to an anaerobic chamber and equilibrated over 3 days. Following oxygen sparging, 55µl of sample was placed on screen printed electrodes (Metrohm) with gold working, gold counter, and silver reference electrodes and analyzed on a chi650e electrochemical workstation.

### Zeta potential

OmcZʹ samples were prepared at 5µM concentration in 5mM Tris + 2mM NaCl at either pH 7.4 or 8.1 and placed into DTS1060 folded capillary cells for zeta potential measurement on zetasizer nanoseries.

### Cryo-EM data collection and processing

OmcZʹ samples were prepared in 0.1X PBS and 20mM ethanolamine pH 10.5 with 2% glycerol. Three-hundred mesh gold quantifoil grids were first glow discharged for 30 s at 20mV operating voltage. Grids were then transferred to an FEI Vitrobot Mark IV operating at 23 °C with 100% humidity followed by 3 µl sample addition to the grids with 3.5 s wait and 5.5 s blotting time before being plunge frozen in liquid ethane.

Frozen grids were imaged on Titan Krios G3 operating at 300 kV equipped with a K3 detector (Gatan). Micrographs were recorded at 0.8332 Å per pixel, with a total dose of ∼70 e^−^ Å^−2^ fractioned into 50 frames. A defocus range of ∼2 µm was used to collect ∼4,700 images for OmcZʹ and ∼8,500 images for OmcZ^+^.

All movies were first motion corrected with MotionCorr and contrast transfer function estimated in CryoSPARC v4.7.2 using all the frames. Filament tracing was used to generate particle libraries originally consisting of ∼2,700,000 for OmcZʹ and ∼4,200,000 for OmcZ^+^ and picked at 360 pixel box size (Fourier cropped to 180px) for initial 2D classification. Following 2D classification, selected particles were re-extracted at 360 pixel box size (Fourier cropped to 300 pixel for OmcZʹ). Selected particles were used in ab-initio reconstruction, followed by heterologous and non-uniform refinement reconstruction, yielding ∼1,101,000 particles for OmcZʹ and ∼356,700 particles for OmcZ^+^. Symmetry search from non-uniform refinement resulted in initial estimates of 57.6Å rise and -160.2° pitch for OmcZʹ and 58.2Å rise and -172.9° pitch for OmcZ+ before further processing in helical refinement for final map generation. Helical refinement in OmcZʹ resulted in a 2.9Å resolution map that was used in model refinement. OmcZ^+^ was further processed using 2D classification to clean dendritic elements from nanowire selections, resulting in ∼190,913 particles used in final helical reconstruction of the 3.4Å map.

Chain B of PDB:7LQ5^3^ was mutated in pymol for both OmcZʹ and OmcZ^+^ before using as the initial model for map fitting in Phenix 2.0-5936. Independent half maps of both OmcZʹ and OmcZ+ were used in Phenix Resolve to generate density modified maps for use in model refinement. Global model refinement was performed using Phenix Real-space Refinement and local refinement performed in Coot. Following refinement, the model was transformed along helical rise and twist parameters from cryoSPARC helical refinement using ChimeraX, then fit to map density to form the representative nanowire. Nanowire refinement was performed locally in Coot and globally in Phenix to generate final models (36HZ / EMD-77591: OmcZʹ, 36KE / EMD-77621: OmcZ^+^).

### Fabrication of OmcZ hydrogel WEG

OmcZʹ samples were concentrated and washed with pure water to remove residual buffer and salts. Nanowire samples were resuspended in 400µL of milliQ water and compared against weighted standards until roughly 8µg of nanowire material was resuspended into solution. The absorbance of these samples was then recorded and used in aiding consistency of future nanowire solutions. Nanowire solutions were added to a 1cm x 2cm x 2mm mold on glass substrate with stir bar on a stir plate at 60 RPM. 8µg of sodium alginate was weighed out and slowly dispensed into the nanowire solution. The stir bar was then removed and the nanowire:alginate solution was suspended into a 1.2% CaCl_2_ bath for 1 hour for matrix crosslinking. Formed hydrogels were removed from the mold and stored on glass substrates atop hydrated filter paper in petri dishes at 4C. Prior to electrical characterization hydrogels were subject to washing in pure water for 4 hours to remove residual ions.

The prepared OmcZʹ hydrogel samples were placed on a glass substrate to provide mechanical support during testing. Carbon cloth electrodes (0.5 cm in width) were attached to both ends of the hydrogel and secured using wooden clips to ensure stable electrical contact. The assembled device was then positioned vertically in a Petri dish, and the working solution required for evaporation-driven electricity generation was added until the liquid level reached and fully covered the lower electrode, while the upper electrode remained exposed to air.

### Hydrogel Rheometry

Rheological characterization was performed using a DHR20 rheometer from TA Instruments, with a 8 mm parallel plate geometry. Gel samples were prepared by trimming to the appropriate size using a punch cutter. All measurements were taken at ambient temperature, and samples were surrounded with mineral oil after loading to prevent evaporation during the measurement. First, to determine the storage modulus (G’) and loss modulus (G”), time sweeps of up to 120 s were recorded at constant strain of 1%, and constant angular frequency of 1 rad/s. Further, to confirm shear thinning behavior, strain sweeps were recorded at a constant angular frequency of 1 rad/s, and G’ and G” were recorded as a function of oscillation strain % from 0.1 to 100. Data was collected using the TRIOS software (n = 2 technical replicates for OmcZ-alginate and n = 1 for alginate hydrogel).

### Electrical measurements

Voltage and current were recorded using a BioLogic VMP3 potentiostat in a two-electrode configuration.

## ACKNOWLEDGEMENTS

We thank the Gralnick Lab (University of Minnesota) for *S. oneidensis* strains JG596 (ϕλ3), JG1194 (ϕλ8), JG1519 (ϕλ10), as well as the Baker Lab (University of Washington) for assistance in phylogenic biomining. The authors would also like to thank the University of Texas at Austin core facilities Sauer Structural Biology Laboratory (RRID:SCR_022951) and Biological Mass Spectrometry Facility (RRID:SCR_021728) for their assistance in data collection. Funding for this work was provided by the Air Force Office of Scientific Research (FA9550-24-1-0273, BKK), National Institutes of Health (R35GM156283, BKK). National Science Foundation (NSF 2227399, ADE), and the Welch Foundation (F-1929 to BKK, F-1654 to ADE, F-1861 to GY). Additional funding was provided by the Army Research Office and accomplished under Cooperative Agreement Number W911NF-22-2-0246. The views and conclusions contained in this document are those of the authors and should not be interpreted as representing the official policies, either expressed or implied, of the Army Research Office or the U.S. Government. The U.S. Government is authorized to reproduce and distribute reprints for Government purposes notwithstanding any copyright notation herein.

## DATA AVAILABILITY

CryoEM analysis, raw images, and structures are available at PDB and EMDB (pending publication) using accession codes 36HZ, EMD-77591 (OmcZ) and 36KE, EMD-7761 (OmcZ^+^). Additional raw data supporting the findings in this manuscript is available through the Texas Data Repository at doi: XXX. Correspondence and requests for materials should be addressed to GY, BKK, and ADE.

## AUTHOR CONTRIBUTIONS

ERS, XL, GY, BKK, and ADE designed the research. ERS, XL, MLR, VD, and LLZ performed the research. AB assisted with CyroEM analysis. ERS, XL, MLR, VD, DW, and LLZ helped analyze data. ERS, BKK, and ADE wrote the paper with input from all authors.

## EXTENDED DATA

**Extended Data Fig. 1.**
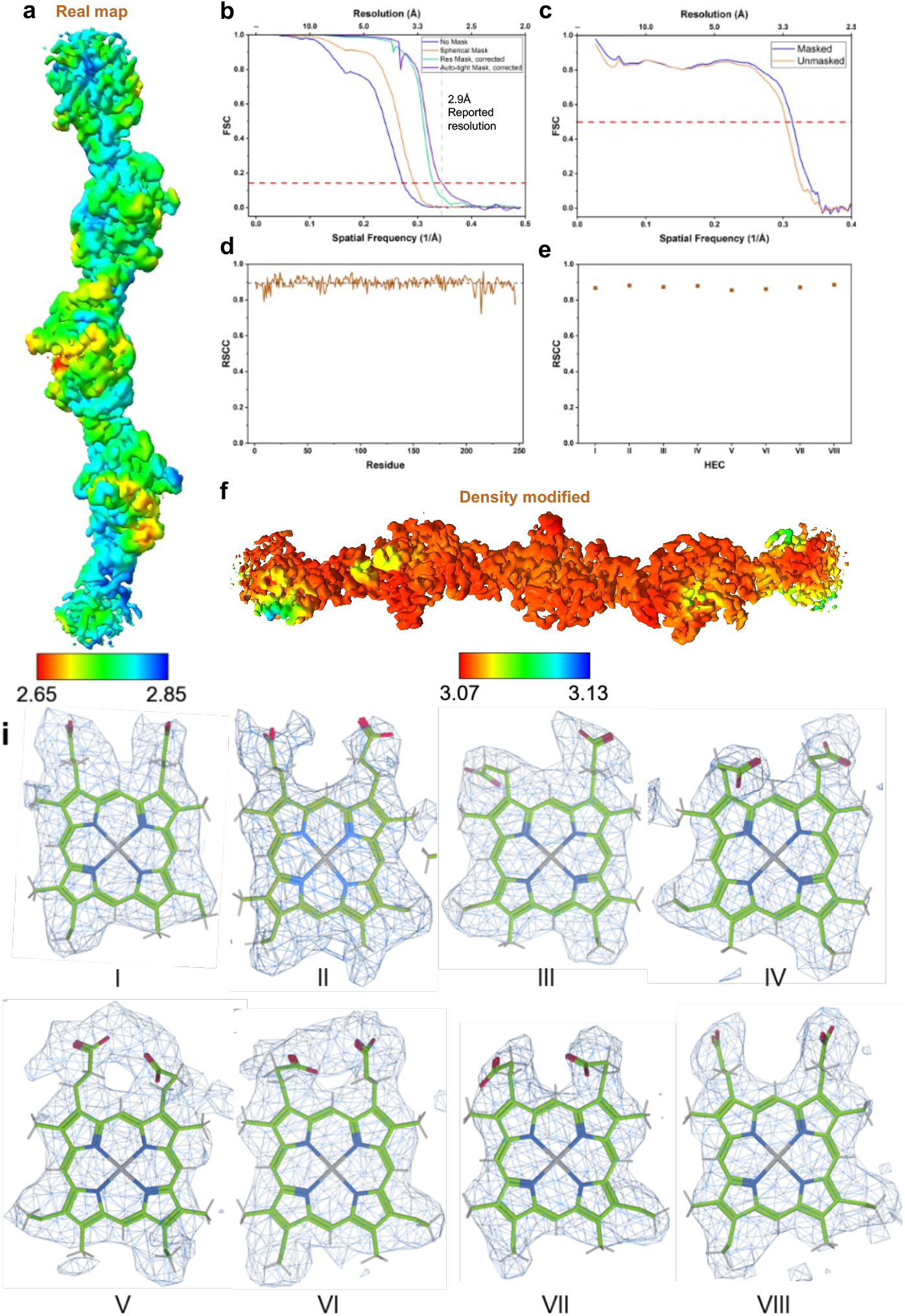
OmcZʹ Cryo-EM map reconstruction and fit quality. (a) Local resolution of half maps displayed on real map generated from cryoSPARC helical reconstruction. (b) Independent half maps FSC with 0.143 gold-standard cut-off (red dash line). (c) Map to model FSC. (d-e) Real space correlation coefficient (RSCC) for residues (left) and hemes (right) using real map generated from cryoSPARC helical refinement. (f) Local resolution of density modified map used in structure refinement. (g) Fitting of heme cofactors at 2.0Å threshold density in map(f).

**Extended Data Fig. 2.**
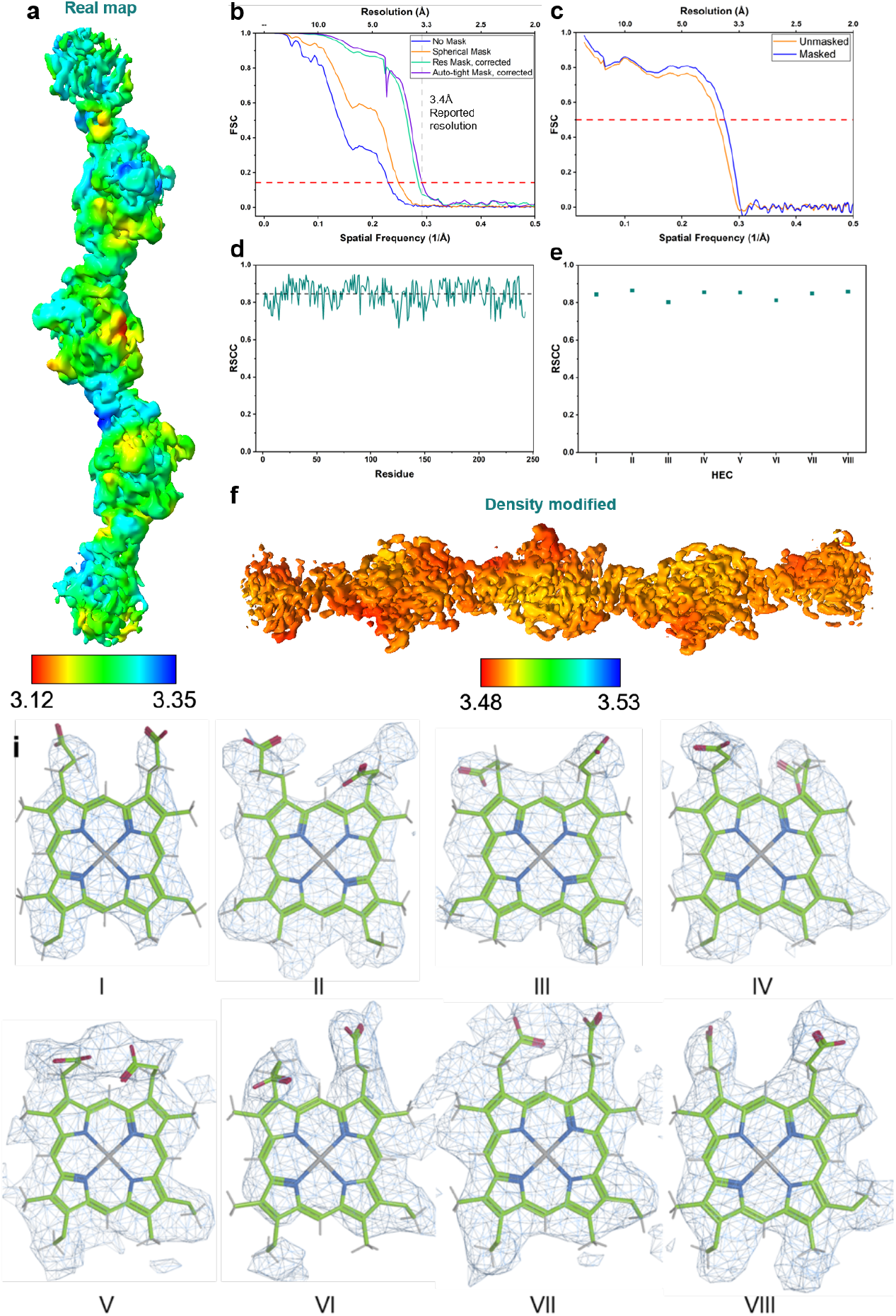
OmcZ^+^ Cryo-EM map reconstruction and fit quality. (a) Local resolution of half maps displayed on real map generated from cryoSPARC helical reconstruction. (b) Independent half maps FSC with 0.143 gold-standard cut-off (red dash line). (c) Map to model FSC. (d-e) Real space correlation coefficient (RSCC) for residues (left) and hemes (right) using real map generated from cryoSPARC helical refinement. (f) Local resolution of density modified map used in structure refinement. (g) Fitting of heme cofactors at 2.0Å threshold density in map(f).

**Extended Data Fig. 3.**
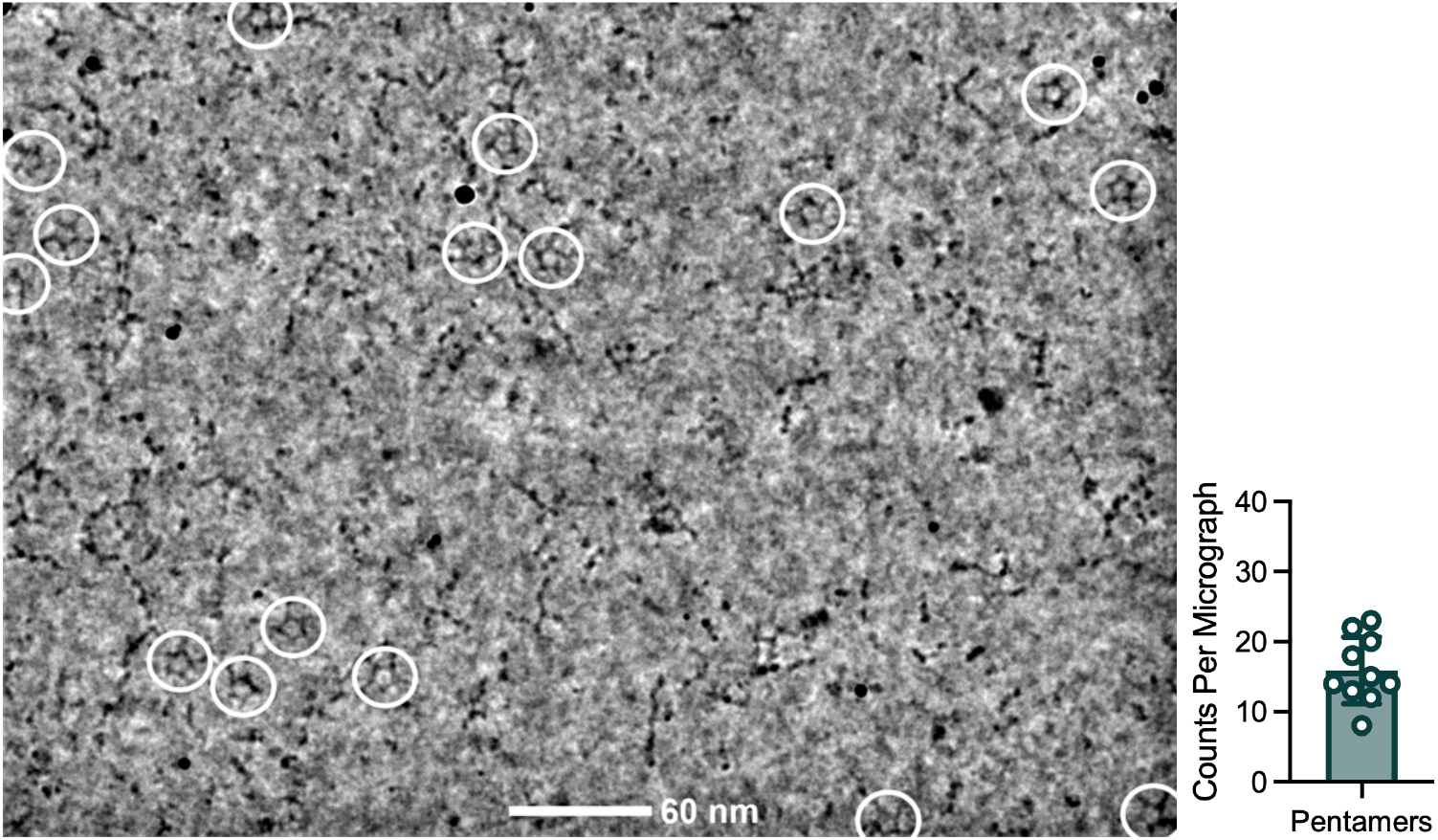
OmcZ^+^ pentamer assembly. (a) OmcZ^+^ micrograph, binned 4, highlighting pentameric assemblies found throughout (white circles). (b) Average count of pentamer assemblies across 10 micrographs.

**Extended Data Fig. 4.**
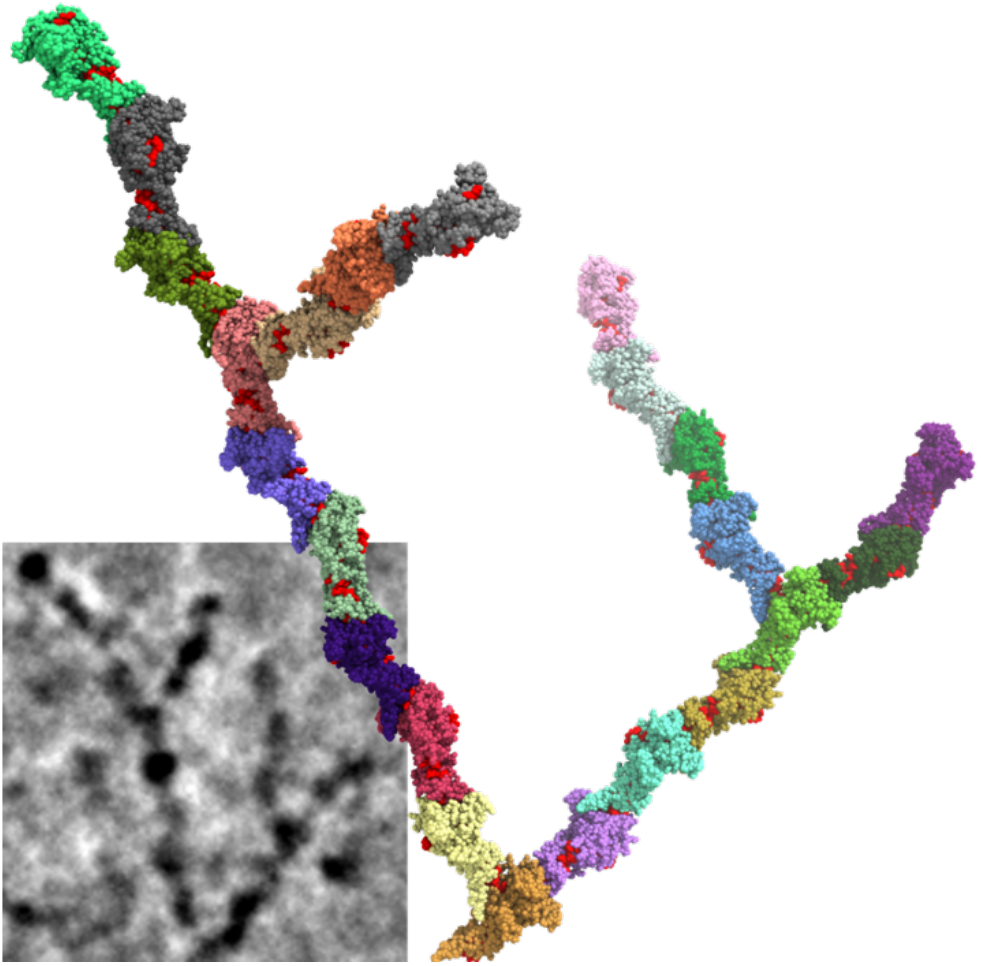
OmcZ^+^ dendritic structural interpretation. Representative cryo-EM micrograph segment (inset) with fitted cryo-EM density segments illustrating the proposed arrangement of OmcZ^+^ protomers within dendritic nanowires. Structural interpretation constructed from OmcZ^+^ nanowire (PDB:36KE) and OmcZ^+^ hemeVII:II dimers (Fig.3h,i) using pymol.

## REFERENCES

1. Yalcin, S.E. et al. Electric field stimulates production of highly conductive microbial OmcZ nanowires. Nat Chem Biol 16, 1136–1142 (2020).

2. Groenendaal, L., Jonas, F., Freitag, D., Pielartzik, H. & Reynolds, J.R. Poly(3,4-ethylenedioxythiophene) and Its Derivatives: Past, Present, and Future. Advanced Materials 12, 481–494 (2000).

3. Gu, Y. et al. Structure of Geobacter cytochrome OmcZ identifies mechanism of nanowire assembly and conductivity. Nature Microbiology 8, 284–298 (2023).

4. Nevin, K.P. et al. Anode Biofilm Transcriptomics Reveals Outer Surface Components Essential for High Density Current Production in Geobacter sulfurreducens Fuel Cells. PLOS ONE 4, e5628 (2009).

5. Richter, H. et al. Cyclic voltammetry of biofilms of wild type and mutant Geobacter sulfurreducens on fuel cell anodes indicates possible roles of OmcB, OmcZ, type IV pili, and protons in extracellular electron transfer. Energy & Environmental Science 2, 506–516 (2009).

6. Arslan, E., Schulz, H., Zucerey, R., Künzler, P. & Thöny-Meyer, L. Overproduction of theBradyrhizobium japonicum c-Type Cytochrome Subunits of thecbb3Oxidase inEscherichia coli. Biochemical and Biophysical Research Communications 251, 744–747 (1998).

7. Meyer, T.E. et al. Identification of 42 possible cytochrome C genes in the Shewanella oneidensis genome and characterization of six soluble cytochromes. Omics 8, 57–77 (2004).

8. Arnold, R.G., Hocmann, M.R., Dichristina, T.J. & Picardal, F.W. Regulation of Dissimilatory Fe(III) Reduction Activity in Shewanella putrefaciens. Appl Environ Microbiol 56, 2811–2817 (1990).

9. Lies, D.P. et al. Shewanella oneidensis MR-1 uses overlapping pathways for iron reduction at a distance and by direct contact under conditions relevant for Biofilms. Appl Environ Microbiol 71, 4414–4426 (2005).

10. Myers, C.R. & Nealson, K.H. Bacterial manganese reduction and growth with manganese oxide as the sole electron acceptor. Science 240, 1319–1321 (1988).

11. Lovley, D.R., Phillips, E.J. & Lonergan, D.J. Hydrogen and Formate Oxidation Coupled to Dissimilatory Reduction of Iron or Manganese by Alteromonas putrefaciens. Appl Environ Microbiol 55, 700–706 (1989).

12. Ruebush, S.S., Brantley, S.L. & Tien, M. Reduction of soluble and insoluble iron forms by membrane fractions of Shewanella oneidensis grown under aerobic and anaerobic conditions. Appl Environ Microbiol 72, 2925–2935 (2006).

13. Edwards, M.J., White, G.F., Butt, J.N., Richardson, D.J. & Clarke, T.A. The Crystal Structure of a Biological Insulated Transmembrane Molecular Wire. Cell 181, 665–673.e610 (2020).

14. Li, Y. et al. Metabolic regulation of Shewanella oneidensis for microbial electrosynthesis: From extracellular to intracellular. Metabolic Engineering 80, 1–11 (2023).

15. Coursolle, D. & Gralnick, J.A. Modularity of the Mtr respiratory pathway of Shewanella oneidensis strain MR-1. Molecular Microbiology 77, 995–1008 (2010).

16. Coursolle, D. & Gralnick, J.A. Reconstruction of Extracellular Respiratory Pathways for Iron(III) Reduction in Shewanella Oneidensis Strain MR-1. Front Microbiol 3, 56 (2012).

17. Edwards, Marcus J. et al. The Crystal Structure of the Extracellular 11-heme Cytochrome UndA Reveals a Conserved 10-heme Motif and Defined Binding Site for Soluble Iron Chelates. Structure 20, 1275–1284 (2012).

18. Thirumurthy, M.A. & Jones, A.K. Geobacter cytochrome OmcZs binds riboflavin: implications for extracellular electron transfer. Nanotechnology 31, 124001 (2020).

19. Inoue, K. et al. Purification and characterization of OmcZ, an outer-surface, octaheme c-type cytochrome essential for optimal current production by Geobacter sulfurreducens. Appl Environ Microbiol 76, 3999–4007 (2010).

20. Beliaev, A.S. et al. Gene and Protein Expression Profiles of Shewanella oneidensis during Anaerobic Growth with Different Electron Acceptors. OMICS: A Journal of Integrative Biology 6, 39–60 (2002).

21. Bhadra, S. et al. Producing molecular biology reagents without purification. PLOS ONE 16, e0252507 (2021).

22. Dong, Z., Guo, S., Fu, H. & Gao, H. Investigation of a spontaneous mutant reveals novel features of iron uptake in Shewanella oneidensis. Scientific Reports 7, 11788 (2017).

23. Dawson, J.H. Probing structure-function relations in heme-containing oxygenases and peroxidases. Science 240, 433–439 (1988).

24. Poulos, T.L. & Kraut, J. The stereochemistry of peroxidase catalysis. J Biol Chem 255, 8199–8205 (1980).

25. Banci, L., Bertini, I., Turano, P., Tien, M. & Kirk, T.K. Proton NMR investigation into the basis for the relatively high redox potential of lignin peroxidase. Proc Natl Acad Sci U S A 88, 6956–6960 (1991).

26. Varadarajan, R., Zewert, T.E., Gray, H.B. & Boxer, S.G. Effects of buried ionizable amino acids on the reduction potential of recombinant myoglobin. Science 243, 69–72 (1989).

27. Bhagi-Damodaran, A., Petrik, I.D., Marshall, N.M., Robinson, H. & Lu, Y. Systematic tuning of heme redox potentials and its effects on O2 reduction rates in a designed oxidase in myoglobin. J Am Chem Soc 136, 11882–11885 (2014).

28. Wang, F. et al. Structure of Geobacter OmcZ filaments suggests extracellular cytochrome polymers evolved independently multiple times. eLife 11, e81551 (2022).

29. Szmuc, E. et al. Engineering Geobacter pili to produce metal:organic filaments. Biosens Bioelectron 222, 114993 (2023).

30. Firer-Sherwood, M., Pulcu, G.S. & Elliott, S.J. Electrochemical interrogations of the Mtr cytochromes from Shewanella: opening a potential window. JBIC Journal of Biological Inorganic Chemistry 13, 849–854 (2008).

31. Xu, J., et al. Sustainable moisture energy. Nature Reviews Materials 9, 722–737 (2024).

32. Zhao, F., Guo, Y., Zhou, X., Shi, W. & Yu, G. Materials for solar-powered water evaporation. Nature Reviews Materials 5, 388–401 (2020).

33. Yao, J. & Song, Y. Calling for sustainable mechanisms in moisture-based hydrovoltaic devices. Natl Sci Rev 12, nwaf315 (2025).

34. Liu, X. et al. Microbial biofilms for electricity generation from water evaporation and power to wearables. Nat Commun 13, 4369 (2022).

35. Liu, X. et al. Power generation from ambient humidity using protein nanowires. Nature 578, 550–554 (2020).

36. Hu, Q., Ma, Y., Ren, G., Zhang, B. & Zhou, S. Water evaporation&#x2013;induced electricity with *Geobacter sulfurreducens* biofilms. Science Advances 8, eabm8047 (2022).

37. Guberman-Pfecer, M.J., Dorval Courchesne, N.M. & Lovley, D.R. Microbial nanowires for sustainable electronics. Nature Reviews Bioengineering 2, 869 – 886 (2024).

38. Xue, G. et al. Water-evaporation-induced electricity with nanostructured carbon materials. Nat Nanotechnol 12, 317–321 (2017).

39. Zhou, X. et al. Harvesting Electricity from Water Evaporation through Microchannels of Natural Wood. ACS Appl Mater Interfaces 12, 11232–11239 (2020).

40. Wang, X. et al. Hydrovoltaic technology: from mechanism to applications. Chemical Society Reviews 51, 4902–4927 (2022).

41. Wang, Z. et al. Unipolar Solution Flow in Calcium-Organic Frameworks for Seawater-Evaporation-Induced Electricity Generation. J Am Chem Soc (2024).

42. Shen, C. et al. A widespread and ancient bacterial machinery assembles cytochrome OmcS nanowires essential for extracellular electron transfer. Cell Chem Biol 32, 239–254.e237 (2025).

43. Petersen, H.A. et al. A Bundled Antiparallel Cytochrome Nanowire Structure Suggests Roles in Cell-Cell Electron Transfer and Biofilm Formation. bioRxiv, 2025.2010.2022.684050 (2025).

44. Cao, M. et al. Ambient-Dried Nanocellulose Composite Aerogels for Enhanced Hydrovoltaic Electricity Generation. Advanced Functional Materials 35, 2418823 (2025).

45. An, N. et al. High-efficiency evaporation induced electricity generation of wood in seawater. Nano Energy 153, 111928 (2026).

46. Lin, J. et al. All Wood-Based Water Evaporation-Induced Electricity Generator. Advanced Functional Materials 34, 2314231 (2024).

47. Anwar, T. & Tagliabue, G. Salinity-dependent interfacial phenomena toward hydrovoltaic device optimization. Device 2, 100287 (2024).

48. Zhang, Z. et al. Integrating Photoelectrochemical Feature on a Hydrovoltaic Chip with High-Salinity Adaption as a Self-Powered Device for Formaldehyde Monitoring. ACS Sensors 9, 2520–2528 (2024).

49. Wang, L., Liu, L. & Solin, N. Ionovoltaic electricity generation over graphene-nanoplatelets: protein-nanofibril hybrid materials. Nanoscale Advances 5, 820–829 (2023).

50. Park, J.H., Park, S.H., Lee, J. & Lee, S.J. Solar Evaporation-Based Energy Harvesting Using a Leaf-Inspired Energy-Harvesting Foam. ACS Sustainable Chemistry & Engineering 9, 5027–5037 (2021).

51. Luo, G., et al. Highly Stretchable, Knittable, Wearable Fiberform Hydrovoltaic Generators Driven by Water Transpiration for Portable Self-Power Supply and Self-Powered Strain Sensor. Small 20, 2306318 (2024).

52. He, N. et al. Ion engines in hydrogels boosting hydrovoltaic electricity generation. Energy & Environmental Science 16, 2494–2504 (2023).

53. Wang, C., Tang, S., Li, B., Fan, J. & Zhou, J. Construction of hierarchical and porous cellulosic wood with high mechanical strength towards directional Evaporation-driven electrical generation. Chemical Engineering Journal 455, 140568 (2023).

