## Supplemental Info for "Expression and Engineering of Conductive Cytochrome Nanowires"

| Name | Reference | Description |
|---|---|---|
| *Shewanella oneidensis* MR-1 | ATCC | Passed lab strain |
| *Shewanella oneidensis* JG596 | D. Coursolle, *et al*. 2010 | MR-1 *ΔmtrCΔomcAΔmtrF* |
| *Shewanella oneidensis* JG1519 | D. Coursolle, *et al*. 2012 | MR-1 *ΔmtrBΔmtrCΔomcAΔmtrFΔmtrA ΔmtrDΔdmsEΔSO4360ΔcctAΔrecA* |
| *Shewanella oneidensis JG1194* | D. Coursolle, *et al*. 2012 | MR-1 *ΔmtrAΔdmsEΔmtrDΔcctA ΔSO_4360ΔmtrCΔmtrFΔOmcA* |
| *Shewanella oneidensis::*Δ4 | E. Szmuc *et al*. 2023 | MR-1 *ΔmshAΔflgKΔpilA + G.s pilA* |
| pESoZgs | This study | *Geobacter sulfurreducens* native OmcZ + OzpA + PI |
| pESoZ' | This study | OmcZ del56-67 + OzpA + PI |
| pESoZ30gfp | This study | OmcZ'-GFP + OzpA + PI |
| pESoZ50gfp | This study | OmcZ'-GFP + PI |
| pESoZsII | This study | OmcZ'-strepII + PI |
| pESoAsII | This study | OzpA-strepII + PI |
| pESoΔPI | This study | OmcZ'-GFP expression only |
| pESoZI13 | This study | OmcZ' [I13E] + OzpA + PI |
| pESoZT82D | This study | OmcZ' [T82D] + OzpA + PI |
| pESoZH85 | This study | OmcZ' [H85M] + OzpA + PI |
| pESoZV120 | This study | OmcZ' [V120D] + OzpA + PI |
| pESoZT173 | This study | OmcZ' [T173D] + OzpA + PI |
| pESoZI13V120 | This study | OmcZ' [I13E,V120D] + OzpA + PI |
| pESoZI13T173 | This study | OmcZ' [I13E,T173D] + OzpA + PI |
| pESoZI13V120T173 | This study | OmcZ' [I13E,V120D,T173D] + OzpA + PI |
| pESoZI13T82V120T173 | This study | OmcZ' [I13E,T82D,V120D,T173D] + OzpA + PI |
| pESoZV7sII | This study | OmcZ'-strepII [V7E] + PI |
| pESoZI13sII | This study | OmcZ'-strepII [I13E] + PI |
| pESoZV37sII | This study | OmcZ'-strepII [V37E] + PI |
| pESoZQ39sII | This study | OmcZ'-strepII [Q39Y] + PI |
| pESoZT82DsII | This study | OmcZ'-strepII [T82D] + PI |
| pESoZT82EsII | This study | OmcZ'-strepII [T82E] + PI |
| pESoZH85MsII | This study | OmcZ'-strepII [H85M] + PI |
| pESoZI103DsII | This study | OmcZ'-strepII [I103D] + PI |
| pESoZS119DsII | This study | OmcZ'-strepII [I103E] + PI |
| pESoZV120DsII | This study | OmcZ'-strepII [V120D] + PI |

| Name | Reference | Description |
|---|---|---|
| pESoZV120TsII | This study | OmcZ'-strepII [V120T] + PI |
| pESoZN166DsII | This study | OmcZ'-strepII [N166D] + PI |
| pESoZQ167EsII | This study | OmcZ'-strepII [N167E] + PI |
| pESoZT169DsII | This study | OmcZ'-strepII [T169D] + PI |
| pESoZT173DsII | This study | OmcZ'-strepII [T173D] + PI |
| pESoZT173EsII | This study | OmcZ'-strepII [T173E] + PI |
| pESoZQ192EsII | This study | OmcZ'-strepII [Q192E] + PI |
| pESoZL202EsII | This study | OmcZ'-strepII [L202E] + PI |
| pESoZI205DsII | This study | OmcZ'-strepII [I205D] + PI |
| pESoZI205EsII | This study | OmcZ'-strepII [I205E] + PI |
| pESoZ5sII | This study | OmcZ'-strepII [I13E,V120D] + PI |
| pESoZ15sII | This study | OmcZ'-strepII [I13E,V120D,T173D] + PI |
| pESoZ36sII | This study | OmcZ'-strepII [I13E,T82D,V120D,T173D] + PI |
| pESoZwt+ | This study | OmcZ-strepII [A42M,N45Q,V47S,T48Y,Δ49-51,N55D,Δ56-67] + PI |
| pESoZ+ | This study | OmcZ-strepII [I13E,A42M,N45Q,V47S,T48Y,Δ49-51,N55D,Δ56-67, T82D,S119D,V120T] + PI |

**Table S1.** Bacterial strains and plasmids used

| Name | Sequence (5' – 3') |
| --- | --- |
| OmcZ[Δ56-67] gBlock | TAATAAGGTCTCATTGTTTAATGATGATTAAAACTATTGCAGGGAAATATCATGAGTAAAAA<br>ACTGCTTTCCGTGCTGTTTGGTGCATCGCTGGCCGCTTTAGCATTGAGCCCGACCGCCTT<br>CGCGGCTGTTCCGCCCCCGCCGGTAAACCAGTTCCTGGGGATCTACGATACCAAGTTTC<br>CCAACCTGACCAAAGCCGACTGTCTCGAGTGTCACGTCAGCGACACGGTGCTCGTCCAG<br>CAGCATCACGCCCTGATCAACACGGTTACGCCGCCGGCAAGCTGCATCAATTGCCACGT<br>CATGGTTCCCGACGGCTCGGGCGGTTTTACCTTCCAGGATTTCAGAAACTGTTTCAACTG<br>TCACACCCAGACTCCGCACCACACCTCTCCCGCGGCTGTTGCCAAGGACTGCAAGTACT<br>GTCACGGCAACTTCATCGACAACCCGCTTGACGGCCACTACATTCCGACCTACAGCGCA<br>AGCTCCGTCACTCCGATGCCGAGCGGTAGGTCTGTAACCGCTACGGATGGCAATGTTGT<br>TATTGTTCAGGGTTGCGAAGCATGCCATCAGGCAGCGCCCAACGCCATCGATCCGAAGA<br>CCAATACTGTCCGCCCGATCTTCAGCAACCAGGATACTCACCACGGCACCGGCATCACC<br>GACTGCAACCTCTGCCACAATACGTCCTCCAATGTACCGATCCGTCAGTGCGAGGTCTGC<br>CACGGTGTGAACTCGCTTCATAATATCCAGAAGGACAGCCCCAATGCAGCCAACCTCGG<br>AACGGTCAAGCCGGGGCTGGAGGATCTGGGCTGGGGCCACATCGGTAACAACTGGGAC<br>TGCCAGGGCTGCCACTGGTCCTGGTTCGGTAATTCCTCGCCGTACACCAATGCCACCGT<br>GCCCGCCATTAACGGCCAGAGCAGCTATACGGTCACCGCGGGCAAGGAAGCCGTCCTG<br>ACCATCGTCGGCTCCAGCTTCGTCAACGTCGGACCCGACGGCGTGACCACCTATCAGCC<br>CACTGTTGCGCTCGTCAGCGGCAGCACGAGCCTGACACTCACTCCCTTCTCGGTCACTG<br>AAAGCGAGATCAAGGTGTCCGTACCCGCCCTCGTTGAAGGTGTCTACGAGCTCCGCATC<br>ACCAAGGCCAACAAGGTCAGCAACCTTGCCAAGCTGACCGTCGCGCCGGCCCGGATCAT<br>TGCCTCCGCTACCCTTGCGACTGGCAAGACCCTTACCATTACCGGCACTGGTTTCGGAC<br>CGGCACCGAGCTCCGAGTACGACGCTGGTATCGGTGTGTACGCCGGAACCACCCAGGC<br>AAACGTCATTTCCTGGAGCGACACCAAGGTGGTTGCCACCAGCCCGGACTTTGCAACCA<br>ACGGCTACGTGACCGTAAAAACCATCAACGGTCCGCTCTCCGGCAAAATCCTGGCGGCT<br>CCGAAGAAAGTCAAACGGTAATGAGACCTTATTA |
| Native OmcZ gBlock | TAATAAGGTCTCATTGTTTAATGATGATTAAAACTATTGCAGGGAAATATCATGAGTAAAAA<br>ACTGCTTTCCGTGCTGTTTGGTGCATCGCTGGCCGCTTTAGCATTGAGCCCGACCGCCTT<br>CGCGGCTGTTCCGCCCCCGCCGGTAAACCAGTTCCTGGGGATCTACGATACCAAGTTTC<br>CCAACCTGACCAAAGCCGACTGTCTCGAGTGTCACGTCAGCGACACGGTGCTCGTCCAG<br>CAGCATCACGCCCTGATCAACACGGTTACGCCGCCGGCAAGCTGCATCAATACCTCCGG<br>CACTGTCCCCCCGACCCTTGCCACCGGTTGCCACGTCATGGTTCCCGACGGCTCGGGC<br>GGTTTTACCTTCCAGGATTTCAGAAACTGTTTCAACTGTCACACCCAGACTCCGCACCACA<br>CCTCTCCCGCGGCTGTTGCCAAGGACTGCAAGTACTGTCACGGCAACTTCATCGACAAC<br>CCGCTTGACGGCCACTACATTCCGACCTACAGCGCAAGCTCCGTCACTCCGATGCCGAG<br>CGGTAGGTCTGTAACCGCTACGGATGGCAATGTTGTTATTGTTCAGGGTTGCGAAGCATG<br>CCATCAGGCAGCGCCCAACGCCATCGATCCGAAGACCAATACTGTCCGCCCGATCTTCA<br>GCAACCAGGATACTCACCACGGCACCGGCATCACCGACTGCAACCTCTGCCACAATACG<br>TCCTCCAATGTACCGATCCGTCAGTGCGAGGTCTGCCACGGTGTGAACTCGCTTCATAAT<br>ATCCAGAAGGACAGCCCCAATGCAGCCAACCTCGGAACGGTCAAGCCGGGGCTGGAGG<br>ATCTGGGCTGGGGCCACATCGGTAACAACTGGGACTGCCAGGGCTGCCACTGGTCCTG<br>GTTCGGTAATTCCTCGCCGTACACCAATGCCACCGTGCCCGCCATTAACGGCCAGAGCA<br>GCTATACGGTCACCGCGGGCAAGGAAGCCGTCCTGACCATCGTCGGCTCCAGCTTCGTC<br>AACGTCGGACCCGACGGCGTGACCACCTATCAGCCCACTGTTGCGCTCGTCAGCGGCA<br>GCACGAGCCTGACACTCACTCCCTTCTCGGTCACTGAAAGCGAGATCAAGGTGTCCGTA<br>CCCGCCCTCGTTGAAGGTGTCTACGAGCTCCGCATCACCAAGGCCAACAAGGTCAGCAA<br>CCTTGCCAAGCTGACCGTCGCGCCGGCCCGGATCATTGCCTCCGCTACCCTTGCGACTG<br>GCAAGACCCTTACCATTACCGGCACTGGTTTCGGACCGGCACCGAGCTCCGAGTACGAC<br>GCTGGTATCGGTGTGTACGCCGGAACCACCCAGGCAAACGTCATTTCCTGGAGCGACAC<br>CAAGGTGGTTGCCACCAGCCCGGACTTTGCAACCAACGGCTACGTGACCGTAAAAACCA<br>TCAACGGTCCGCTCTCCGGCAAAATCCTGGCGGCTCCGAAGAAAGTCAAACGGTAATGA<br>GACCTTATTA |

| Name | Sequence (5' – 3') |
|---|---|
| OzpA gBlock | TAATAAGGTCTCAGTAAATGATTAACACTAAATCTAACATGGAAGAAAAATATGAGAATCG<br>ATAAAACCACAAAGATAGCGCTAAGTTTATCCAGCTTAGCGATTGCAATCTCGGCTGGTG<br>TGAATGCGGCCGACAGGAAGGTCATTGTAGGATTTCGCTCCACGGTGGAAAAAAGGGAC<br>GTTCGCCACAAAGAGAAGGTCTATCGGCACGGCGGTCGCGTCAAGAGGACGCACTCTGC<br>GGTAAACGCCATATCGGCAACCCTGTCCGAAGAGGAAATTGAACGCCTGAAGAAAGATC<br>CCGACGTTGCCTACGTGGAGACGGACTTCGTGCTTTCGTCCATTGAACCAGCAGCGGCT<br>TCACCGGAAGAATATGCAGCAGCGTGGGGTGCGCAGCATATCGGTGCAGACCAGGTTGC<br>TGCGGCAGGTATCACTGGTGCTGGTGTTCGTGTTGCAGTTCTCGATACGGGCATTGATTA<br>CACGCATCCCGATTTGAAGGACAACTACAAGGGGGGGTACAACTTTGTAGCAGACAACAA<br>CGATCCCATGGACGATGCATACTCTCTTAGCCATGGCACCCATGTTGCAGGTATCATCGC<br>CGCACGCAACAACGGTACCGGTGTGGTTGGTGTTGCGCCCGCAGCGGAACTCTATGCAG<br>TCAAGGTTCTTAACGGCGGCCTCGGCGGAGAGTTGAGCGACATTATCGCCGGTATCGAG<br>TGGGCCATCGAGAACCGGATGCAGGTCGTCAACATGAGCTTCGGCAGCATGGAGTTCTC<br>CCAGGCGCTCAAGGATGTCTGCGATCTGGCCTATCGATCGGGAATCGTGCTGGTGGCTT<br>CTGCCGGCAATTTCTCGCCGGGGGCCGTACTCTATCCCGCCGCTTTCGATTCGGTCGTG<br>GCGGTTTCCGCCACCTACCAGGACGACACGCTTGGAACGTTTTCCAGTTACGGTCCCCA<br>GGTCGAATTGGCCGCACCGGGGCACAATATCTATTCCACGGCAATCGGTGGCGGTTACC<br>GCATCAACTTCGGCACATCGCAGGCAGCACCCCATGTCACCGGTGCGGCGGCACTTCTC<br>ATCTCTGCGGGCACTACCGACACCAACGGTAACCGTTCCGTTGCAGACGAGGTCAGGCA<br>ACGACTTGCGGCAGCCGCTCGCGACCTGGGTGAAATGGGTAGGGACATCTACTATGGTT<br>ACGGCCTCGTTGACGTAGCCAAGGCCGTTCTGTCGCCGCCGAACATCGAGACGGTGGTC<br>ACCACGCCGCGGGGGAAACGGTGTGCATCTGCTGCAGCCCTTGATCTGGCGAACTCGA<br>CCTACCGGCTGGACATTACGGGAGCGACGTTGCAGGCGCTTGAAGTCCGCGTCGGGAG<br>CGCCGACGGGCCTCTTGTGAGCTTTATCCGCTTCCGGCGTGGGACTGAAGGGGCGGTAT<br>CGTTCAGCTACACGGCATCTGGCACTGTCAGGCTGGTGCTGATCCCCCACGGTAAACCG<br>GGAACATCGGCGCGGGTGACGGCCGTTCCGGAGCAGCTGTAATGATGTGAGACCTTATT<br>A |
| Prolyl Isomerase gBlock | TAATAAGGTCTCAGATGATTAAAACTATTGCAGGGAAATATCATGAAGACAGTACGCAACC<br>TCATTATTACCATTCTGTGCTTGCTGGGAAGTGTTCCGTCAGCACCAGGCGCAGAGACGG<br>CCGTCGCTCCATCCGGCAGAGCGGCTGCGGCAGTTGTGAACGGTGCTGTCATCTTCCGT<br>GACGAACTCGATCATTTCGTCGAATTTGCCTTGAGTCGACGGAGGGGAGCCGGGCGTAA<br>GGTTACCGACGAGCAGAAGAAGAGAGTCGAGCGCCAGGAGCTCGATAAGCTTATTGCGA<br>TGGAGTTGCTGGCCCAGGCGGGCGAAAAGCTGAATCCTGCCGATCTGTCCAAAAAGATT<br>CAGGCCCGTCGTGAAGCAATCGGTTCTAGCGTATCTCGGGGTGGTAGCGTGGTTGCACC<br>GTCCGAGGACAGACTTAACGACACTGCTCGCCGTGACGTCCTTGTGGACGCATATCTGG<br>CCTCTCGCGGTATCGATGCGATCAAGGTTCCAGAAACGGATCTCAGGGCCTATTACGAAA<br>AGAACAAGTCCGGTTTTAAAAAGCCTGAGACAATCGCTGTCCGGCATATCCTGGTGAAGG<br>TCGAAAAAGAGGCTTCCCCGGAGACTCAGGCGGAGGCCCGGAAAAAGATCGAGGGCAT<br>TCGCGACCGGATTGGCGCCGGCGCCGATTTTGCCGTTCTTGCCAGCGAGAGCTCCGATT<br>GCGCCAGTGCCGCCAAGGGCGGAGATCTGGGAGAGATTCAGCGGGGCTTCATGCCGCG<br>TGAGTTCGACCAGGTCGCCTTTTCCCTCAAACCGGGTGAGACGAGCGGAATCGTCAAGA<br>CGCACCATGGGTTTCACATTATCAGGGTGATGGAACGGCATCCCGAAACGGTCAGGACC<br>TTCGAAGAGATGCGGGATTTCATCGAGCAGTATCTGGCAAAGGATTACCAACGCAAGAAG<br>GTCGAGGAAATCGTTGAAGAGCTGAAGCGCGCAGCAACAATAGATATTCGTATTCAATAA<br>CCAGGCTGAGACCTTATTA |

**Table S2.** OmcZ operon gene block sequences

| Mutation | Forward primer (5' – 3') | Reverse primer (5' – 3') |
|---|---|---|
| V7E | AAGTTAGGTCTCAGGAAAACCAGTTCCTGGGGATCTACGATAC | AACTTAGGTCTCTTTCCGGCGGGGGCGGAAC |
| I13E | TAATCTGGTCTCAGGAATACGATACCAAGTTTCCCAACCTG | ATACTTGGTCTCATTCCCCCAGGAACTGGTTTACCG |
| V37E | ACAATTGGTCTCAGAACAGCAGCATCACGCCCTG | TAAGTTGGTCTCAGTTCGAGCACCGTGTCGCTGAC |
| Q39Y | TTCTAAGGTCTCATACCATCACGCCCTGATCAACACG | TTGAAAGGTCTCAGGTACTGGACGAGCACCGTGTC |
| T82D | TAGAATGGTCTCAGACCCGCACCACACCTCTCC | TTACAAGGTCTCAGGTCCTGGGTGTGACAGTTGAAACAGT |
| T82E | ATGTAAGGTCTCAGAACCGCACCACACCTCTCCC | ATTGAAGGTCTCAGTTCCTGGGTGTGACAGTTGAAACAGT |
| H85M | TTGATTGGTCTCACATGACCTCTCCCGCGGCTGTT | ATCAAAGGTCTCACATGTGCGGAGTCTGGGTGTGACAG |
| I103D | AAGTAAGGTCTCAGACGACAACCCGCTTGACGGC | ATCTTAGGTCTCACGTCGAAGTTGCCGTGACAGTACTTGC |
| I103E | AATTCAGGTCTCAGAAGACAACCCGCTTGACGGC | ATGCATGGTCTCACTTCGAAGTTGCCGTGACAGTACTTGC |
| S119D | ATGTAAGGTCTCAGACGTCACTCCGATGCCGAGC | ACTATAGGTCTCACGTCGCTTGCGCTGTAGGTCGG |
| V120D | AGTATAGGTCTCAGACACTCCGATGCCGAGCGG | ATGTAAGGTCTCATGTCGGAGCTTGCGCTGTAGGTCG |
| V120T | AGTATAGGTCTCAACCACTCCGATGCCGAGCGG | ATGTAAGGTCTCATGGTGGAGCTTGCGCTGTAGGTCG |
| N166D | TCAATAGGTCTCAGACCAGGATACTCACCACGGCAC | ATTCAAGGTCTCAGGTCGCTGAAGATCGGGCGGAC |
| Q167E | AGATTAGGTCTCAGAAGATACTCACCACGGCACCGG | AAGTTAGGTCTCACTTCGTTGCTGAAGATCGGGCGGA |
| T169D | ATTCAAGGTCTCAGACCACCACGGCACCGGCATC | ATCTTAGGTCTCAGGTCATCCTGGTTGCTGAAGATCGGG |
| T173D | ATAGATGGTCTCAGACGGCATCACCGACTGCAACC | AATTCAGGTCTCACGTCGCCGTGGTGAGTATCCTGGT |
| T173E | TATCAAGGTCTCAGAAGGCATCACCGACTGCAACC | TGTAATGGTCTCACTTCGCCGTGGTGAGTATCCTGGTT |
| Q192E | ATTCAAGGTCTCAGAATGCGAGGTCTGCCACGG | TAGTAAGGTCTCAATTCACGGATCGGTACATTGGAGGAC |
| L202E | AATGAAGGTCTCAGAACATAATATCCAGAAGGACAGCCCC | TCTATAGGTCTCAGTTCCGAGTTCACACCGTGGCA |
| I205D | TTCTAAGGTCTCAGACCAGAAGGACAGCCCCAATG | TTCTAAGGTCTCAGGTCATTATGAAGCGAGTTCACACCGT |
| I205E | AATGAAGGTCTCAGAACAGAAGGACAGCCCCAATGC | TTGAATGGTCTCAGTTCATTATGAAGCGAGTTCACACCGTG |

| Mutation | Forward primer (5' – 3') | Reverse primer (5' – 3') |
|---|---|---|
| MethA | AACTTAGGTCTCACAGCTATAGTTGC ATCGATTGCCACGTCATGGTTCCC | TGAATTGGTCTCAGCTGGTTTGAATTAAC ATGTGATGCTGCTGGACGAGC |
| CXXCH | TGTAATGGTCTCGCATCAATTGCCAC GTCATGGTTCCCGA | ACAATTGGTCTCTGATGCAGCTTGCCGG CG |
| GFP bb | GTTCAAGGTCTCAGGGTTCTGCTGGT TCTGCTGCTG | CAAATTGGTCTCTAATCTTATTTGTACAGT TCATCCATACCATGCGTG |
| GFP ins | ATATGTGGTCTCACCGTTTGACTTTCT TCGGAGC | ATATGTGGTCTCACCGTTTGACTTTCTTC GGAGC |
| GFP mut bb | ATATGTGGTCTCAGGTTCTGCTGGTT CTGCT | ATATGTGGTCTCTTACGAGCCGATGATTA ATTGT |
| GFP mut ins | ATATGTGGTCTCTCGTATAATGTGTG GAATTGTGA | ATATGTGGTCTCAAACCCCGTTTGACTTT CTT |

**Table S3**. List of primers used

| No. | Reference | Material / Device | Electrode / Collector | DI water / Pure water data | Saline condition and data |
|---|---|---|---|---|---|
| 1 | Cao, M., Zhu, J., Miao, G. et al. Ambient-dried nanocellulose composite aerogels for enhanced hydrovoltaic electricity generation. Adv. Funct. Mater. 35, 2418823 (2025).[44] | Ambient-dried nanocellulose/CNT composite aerogels (ADA) | Aluminum foils | 0.57 μW cm$^{-2}$ | 3.82 μW cm$^{-2}$ (saturated NaCl solution) |
| 2 | An, N. et al. High-efficiency evaporation induced electricity generation of wood in seawater. Nano Energy 153, 111928 (2026).[45] | Sulfonated natural maple wood | carbon electrodes | 6.2 μW cm$^{-2}$ | 15.5 μW cm$^{-2}$ (seawater) |
| 3 | Lin, J. et al. All wood-based water evaporation-induced electricity generator. Adv. Funct. Mater. 34, 2314231 (2024).[46] | Delignified / cellulosic balsa wood WEIG (DBWG / CBWG) | PET meshes coated with conductive carbon paste and copper meshes | 0.0575 μW cm$^{-2}$ | 0.522 μW cm$^{-2}$ (1.2 M $CaCl_2$) |
| 4 | Anwar, T. & Tagliabue, G. Salinity-dependent interfacial phenomena toward hydrovoltaic device optimization. Device 2, 100287 (2024).[47] | Ordered silicon nanopillar | Al and Ag/AgCl | 1 μW cm$^{-2}$ | 8 μW cm$^{-2}$ (0.1 M saline water) |
| 5 | Zhang, Z. et al. Integrating photoelectrochemical feature on a hydrovoltaic chip with high-salinity adaption as a self-powered device for formaldehyde monitoring. ACS Sens. 9, 2520–2528 (2024).[48] | $NH_2$-MIL-125/$TiO_2$ nanotube | Flexible porous fabric electrode coated with PEDOT:PSS and rGO | 1.4 μW cm$^{-2}$ | 0.54 μW cm$^{-2}$ (5 M NaCl) |
| 6 | Wang, L. *et al.* Ionovoltaic electricity generation over graphene-nanoplatelets: protein-nanofibril hybrid materials. *Nanoscale Adv.* 5, 820–829 (2023[49] | GNP:PNF:$AlCl_3$ | conductive carbon paste | 0.013 μW cm$^{-2}$ | 0.014 μW cm$^{-2}$ (1 M NaCl) |
| 7 | Park, J. H. et al. Solar evaporation-based energy harvesting using a leaf-inspired energy-harvesting foam. ACS Sustain. Chem. Eng. 9, 5027–5037 (2021).[50] | Polydimethylsiloxane with macro- and microporous structures mixed with conductive carbon materials | Cu mesh | 1 μW cm$^{-2}$ | 1.6 μW cm$^{-2}$ ( 0.1 M NaCl) |
| 8 | Luo, G. et al. Highly stretchable, knittable, wearable fiberform hydrovoltaic generators driven by water transpiration for portable self-power supply and self-powered strain sensor. Small 20, 2306318 (2024).[51] | o-MWCNTs/o-CB@MS | Conductive silver glue | 4.5 μW cm$^{-2}$ | 15.3 μW cm$^{-2}$ (1 M NaCl) |
| 9 | Xue, G. *et al.* Water-evaporation-induced electricity with nanostructured carbon materials. *Nat. Nanotechnol.* 12, 317–321 (2017).[38] | Carbon black | CNT | 0.053 μW cm$^{-2}$ | ~0 μW cm$^{-2}$ (0.1 M NaCl) |
| 10 | Liu, X. *et al.* Microbial biofilms for electricity generation from water evaporation and power to wearables. *Nat. Commun.* 13, 4369 (2022).[34] | G. sulfurreducens biofilm | Gold | 1 μW cm$^{-2}$ | 0.42 μW cm$^{-2}$ (seawater) |
| 11 | Wang, Z. *et al.* Unipolar solution flow in calcium–organic frameworks for seawater-evaporation-induced electricity generation. *J. Am. Chem. Soc.* 146, 1690–1700 (2024).[41] | Calcium–Organic Frameworks | Cu | 0.027 μW cm$^{-2}$ | 0.31 μW cm$^{-2}$ (seawater) |
| 12 | He, N. *et al.* Ion engines in hydrogels boosting hydrovoltaic electricity generation. *Energy Environ. Sci.* 16, 2494–2504 (2023).[52] | NaAMPS and Aam | Gaphite and Cu | 3 μW cm$^{-2}$ | 5 μW cm$^{-2}$ (seawater) |

| | | | | | |
|---|---|---|---|---|---|
| 13 | Wang, C. et al. Construction of hierarchical and porous cellulosic wood with high mechanical strength towards directional evaporation-driven electrical generation. Chem. Eng. J. 455, 140568 (2023).[53] | Wood | Cu | 0.57 μW $cm^{-2}$ | 4.18 μW $cm^{-2}$ (saturated NaCl) |
| 14 | Our work | OmcZ:alginate hydrogels | Carbon cloth | 8.7 μW $cm^{-2}$ | 25 μW $cm^{-2}$(seawater) |

**Table S4**. References and data values used for Figure 4h

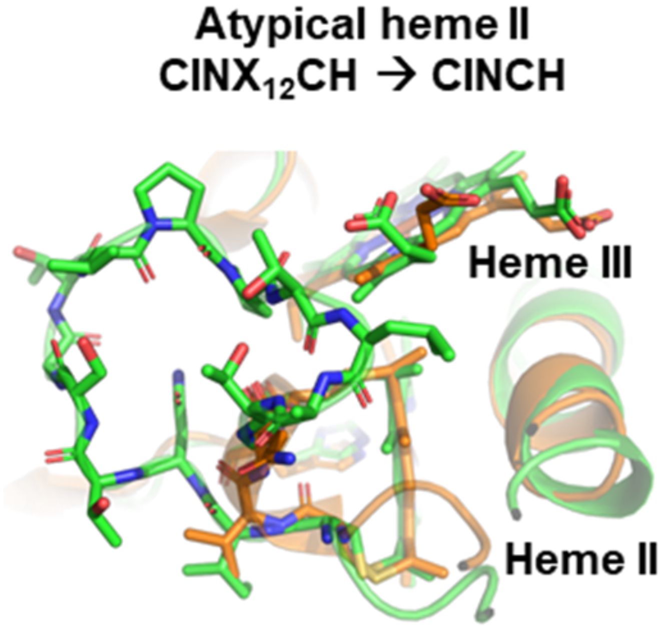


**Fig.S1 *Shewanella* OmcZ' alignment with native *Geobacter* OmcZ.** Alignment of *G sulfurreducens* OmcZ heme II atypical motif CINTSGTVPPTLANKGCH (PDB:8D9M, green) with mutated typical heme motif CINCH of OmcZ' (predicted, orange).

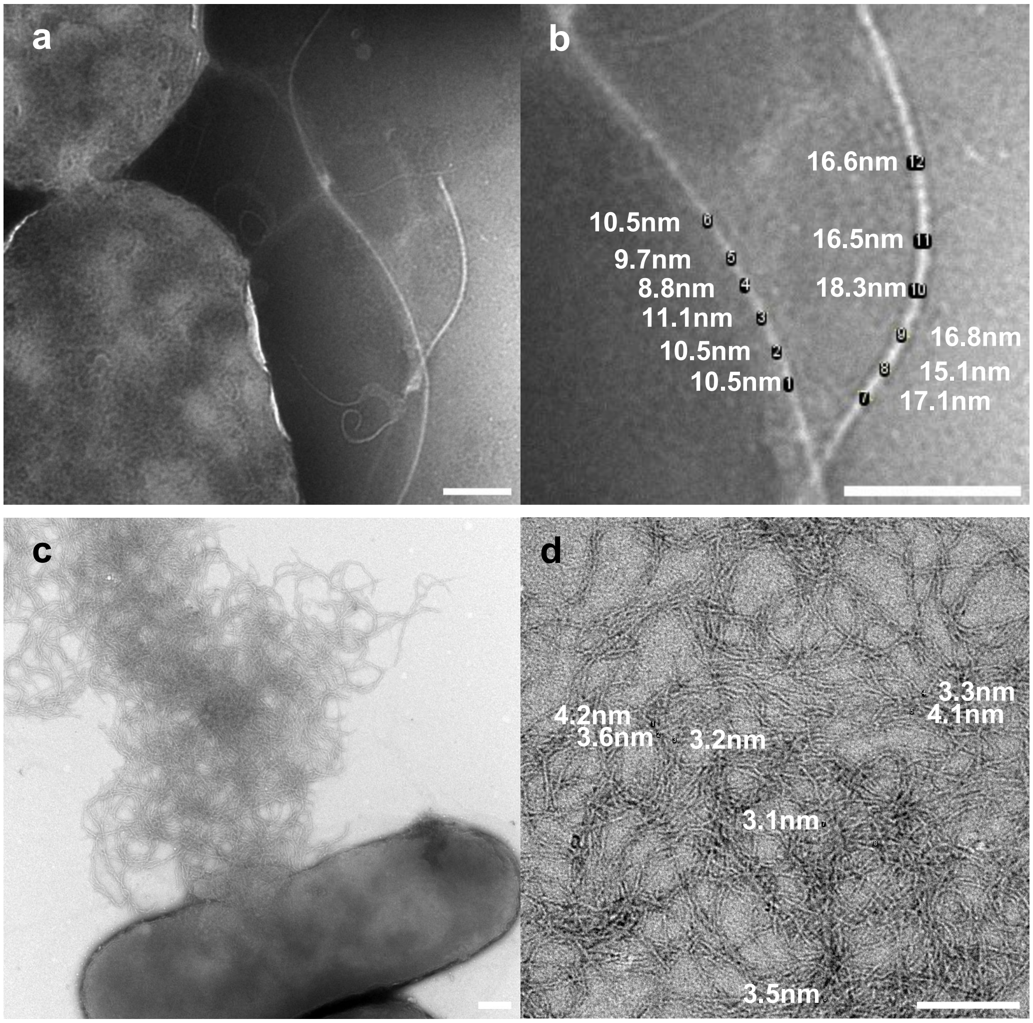


**Fig.S2 *Shewanella* filament size and morphology.** (a) Shewanella MR-1 pili and flagellum under 1% PTA staining. (b) Diameter measurements of pili (left) and flagellum (right). (c) OmcZ' expression in *Shewanella* under 1% PTA staining. (d) Diameter measurements of nanowires from (c). All scale bars 200nm. Measurements taken with ImageJ.

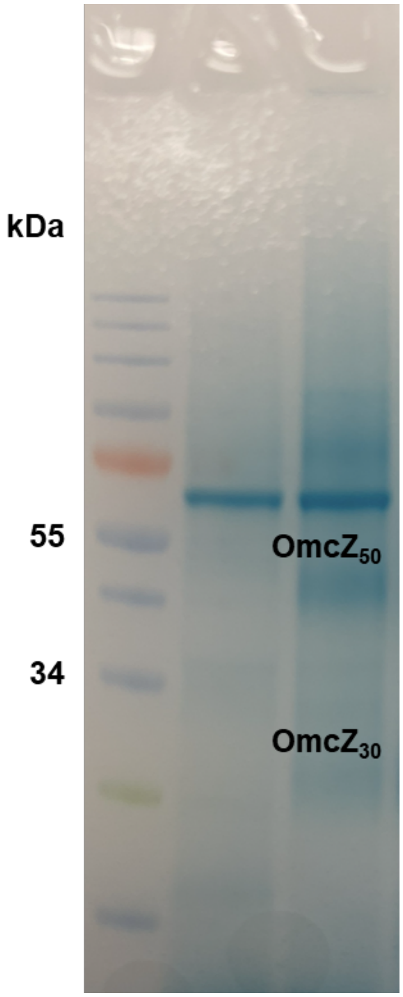


**Fig.S3 Heme stain of *in vivo* OmcZ' expression.** Heme stain of supernatants from *Shewanella oneidensis* MR-1 (lane 1) and same strain expressing OmcZ' plasmid (lane 2).

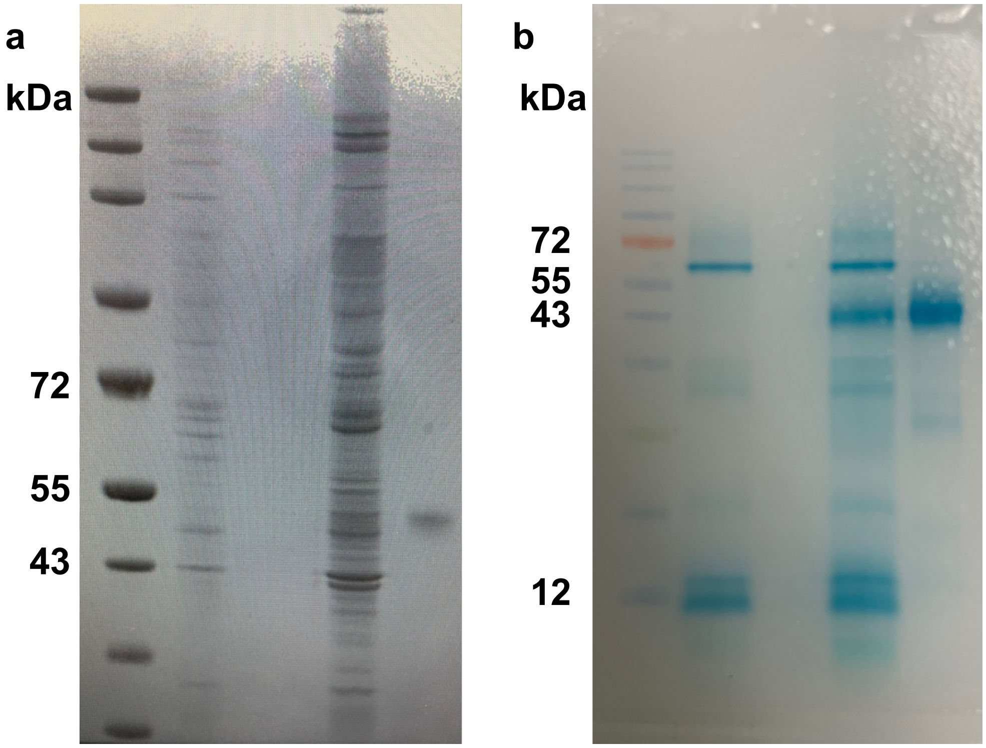


**Fig.S4 *Shewanella* expression of OzpA and OmcZ'$_{50}$.** (a) Coomassie staining of culture supernatants: OzpA utilizing sapSH N-terminal signal sequence pre/post strepII purification (lane1-2), OmcZ'$_{50}$ utilizing cctA N-terminal signal sequence pre/post strepII purification (lane 3-4). (b) Heme staining of culture supernatants: OzpA utilizing sapSH N-terminal signal sequence pre/post strepII purification (lane1-2), OmcZ'$_{50}$ utilizing cctA N-terminal signal sequence pre/post strepII purification (lane 3-4).

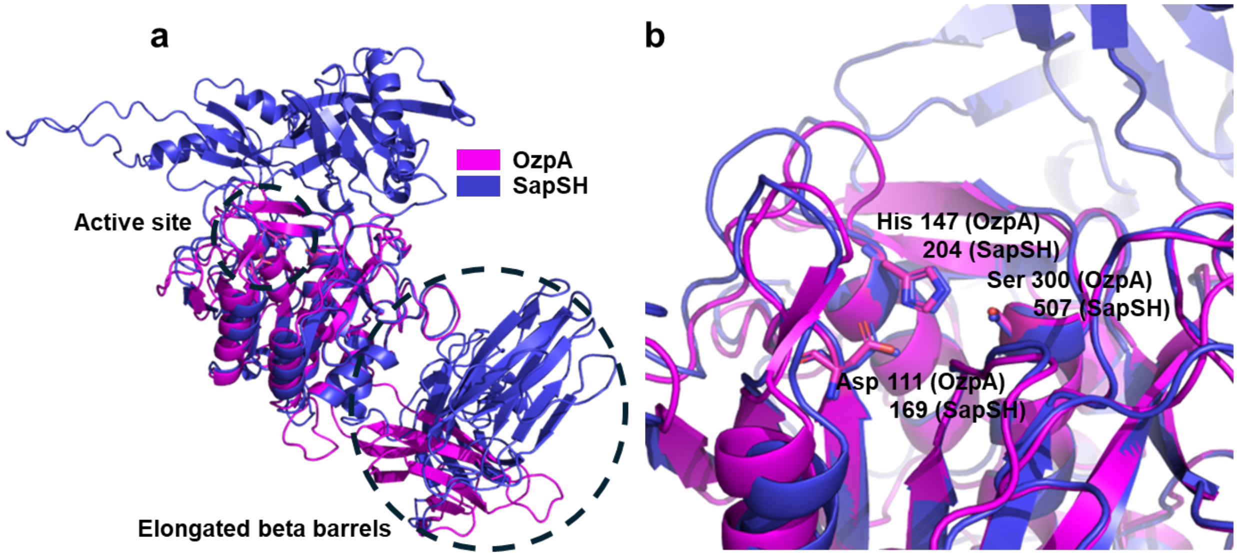


**Fig.S5 Predicted structure comparison of OzpA and SapSH** (a) Structural overlay of subtilases OzpA (magenta) and SapSH (purple). (b) Subtilase active site residues

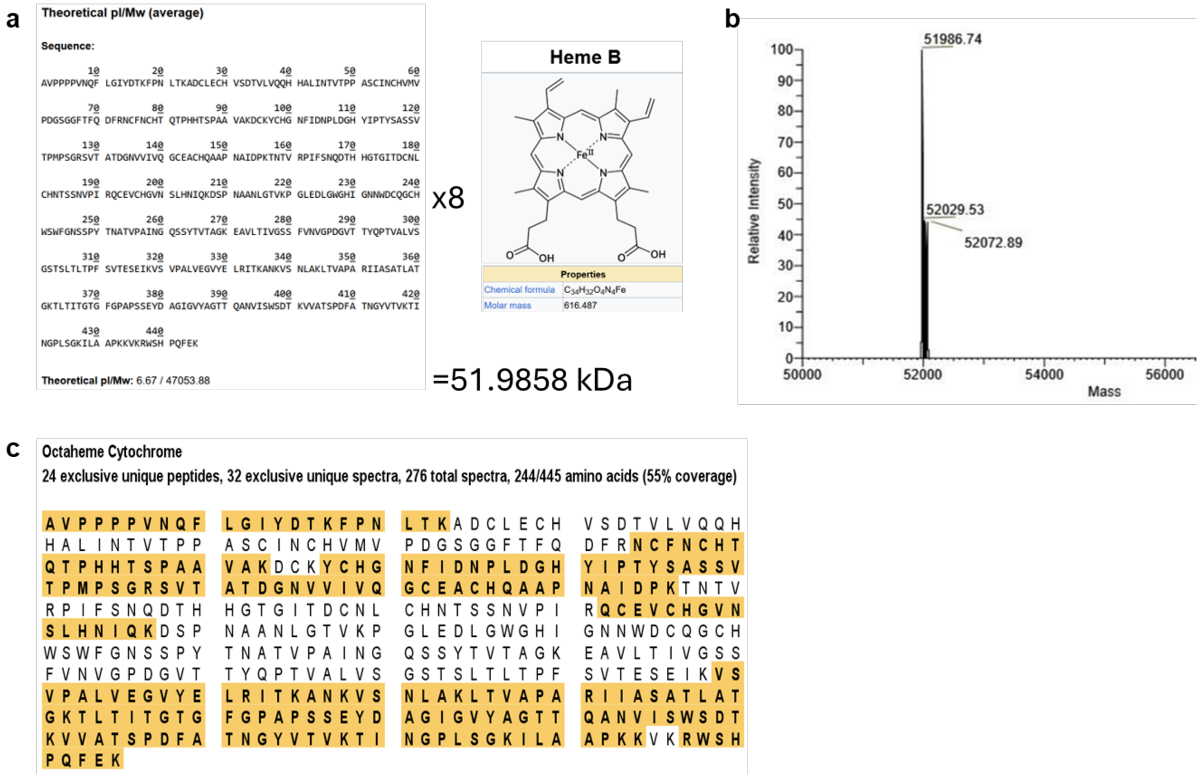


**Fig.S6 StrepII tag OmcZ'$_{50}$ characterization.** (a) Theoretical weight of strepII OmcZ'$_{50}$. (b) Determination of strepII OmcZ'$_{50}$ weight by mass spectroscopy. (c) Liquid Chromatography/ Mass Spectroscopy of strepII OmcZ'$_{50}$ following trypsin digest.

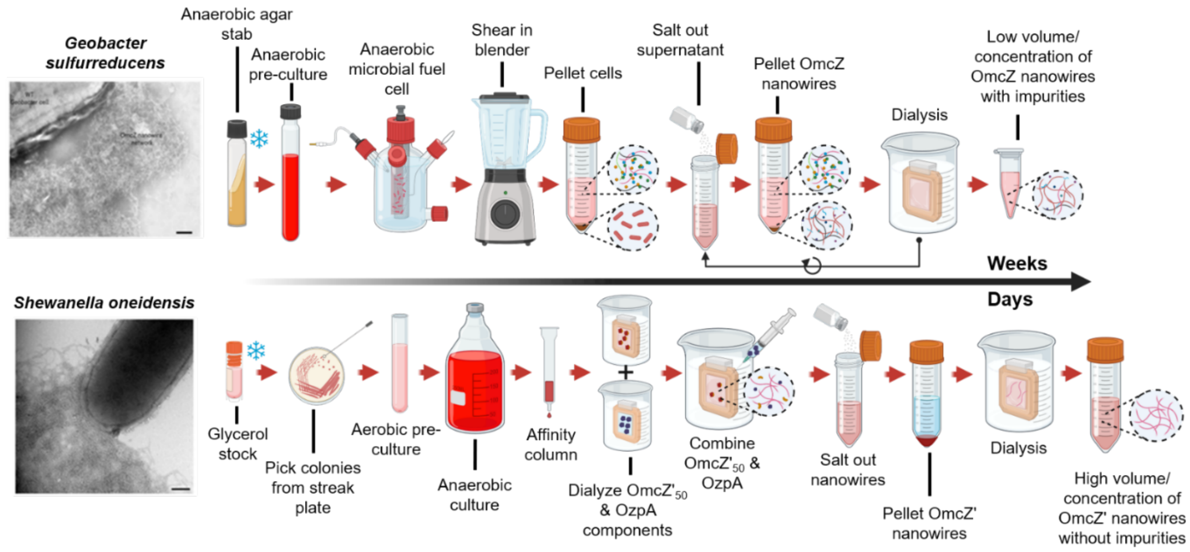


**Fig.S7 Scalable nanowire purification.** Schematic of OmcZ' nanowire *in vitro* purification in *Shewanella* vs *in vivo* nanowire purification from *Geobacter*.

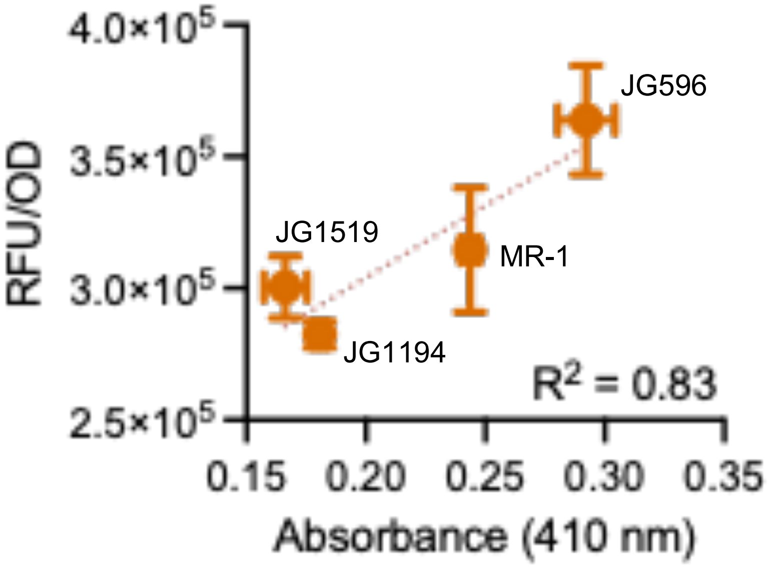


**Fig.S8 Supernatant OmcZ'-GFP absorbance.** 410nm absorbance of *Shewanella* strains MR-1, JG596, JG1194, and JG1519 expressing OmcZ’-GFP.

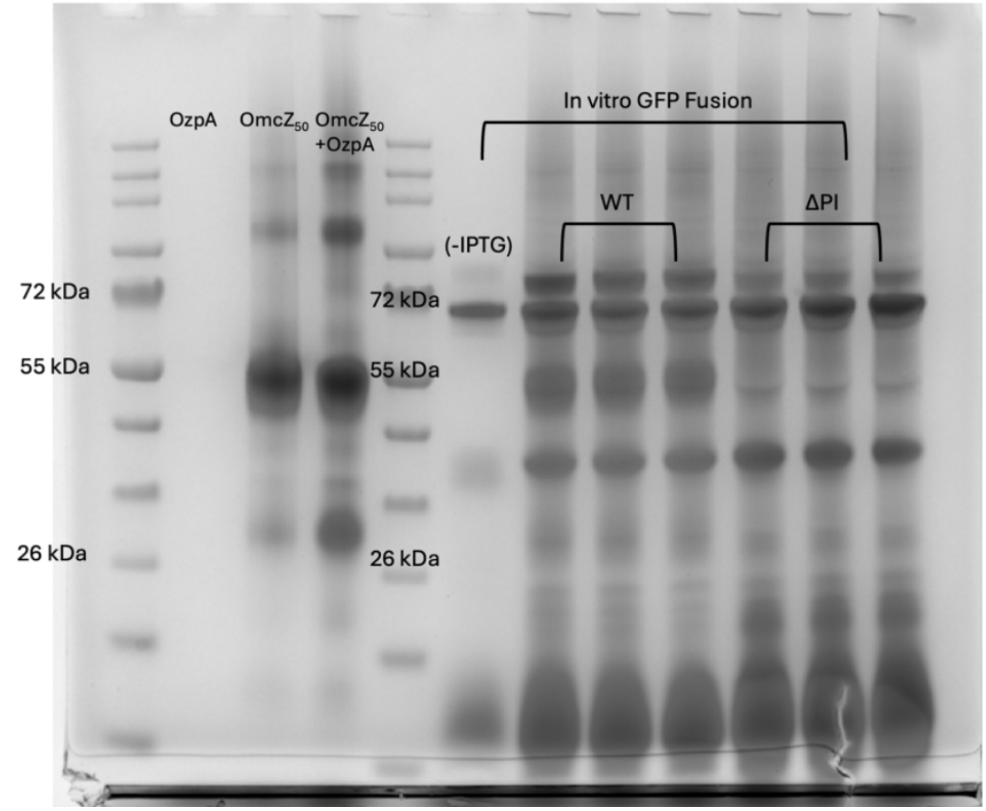


**Fig.S9 Prolyl isomerase in OmcZ' expression.** Heme stain comparison of supernatants from OmcZ'-GFP expression in presence (WT) vs absence (ΔPI) of prolyl isomerase (PI).

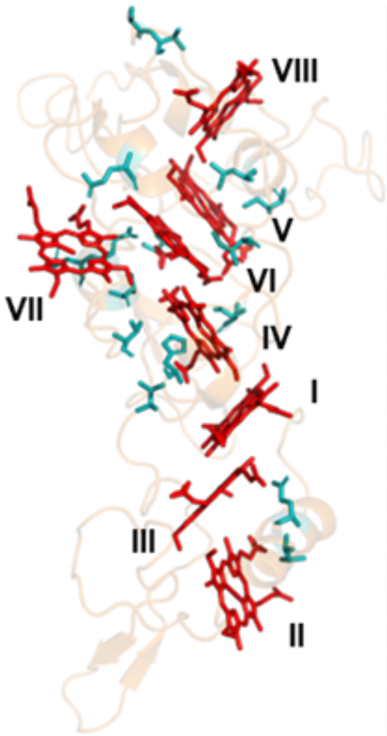


| Mutation | Expression |
|---|---|
| V7E | − |
| I13E | + |
| V37E | − |
| Q39Y | + |
| T82D | + |
| T82E | + |
| H85M | + |
| I103D | − |
| I103E | − |
| S119D | + |
| V120T | + |
| V120D | + |
| N166D | + |
| Q167E | + |
| T169D | + |
| T173D | + |
| T173E | + |
| Q192E | + |
| L202E | − |
| I205D | − |
| I205E | − |

**Fig.S10 Rational mutations in OmcZ'.** Left, native OmcZ (PDB:8D9M, orange) with heme array (red) and rational mutations (teal). Right, table of point mutations indicating whether OmcZ' was expressed (+) or not identified in culture media (-).

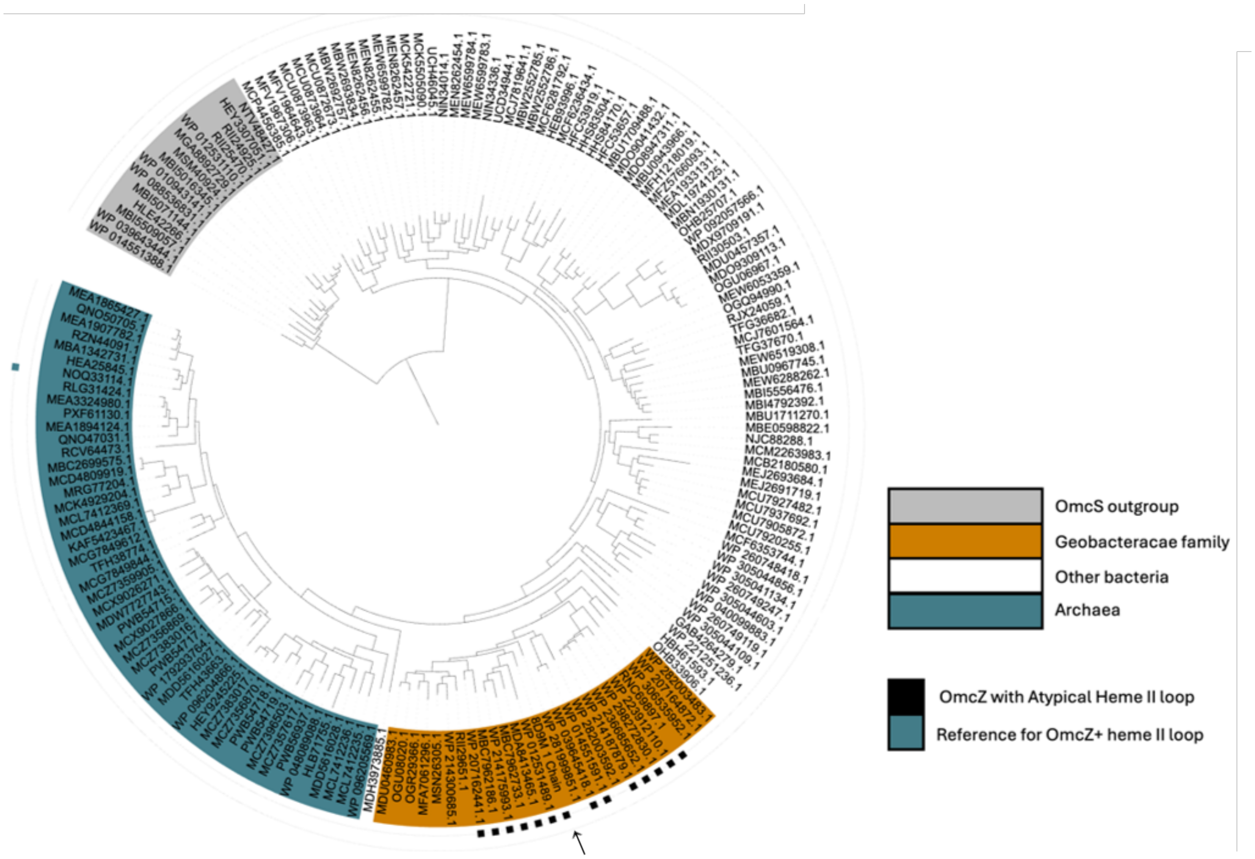


**Fig.S11 Iqtree of NCBI sequence similarity**. Sequence data with 90% cluster identity to *Geobacter sulfurreducens* OmcZ (8D9M) and OmcS outgroup.

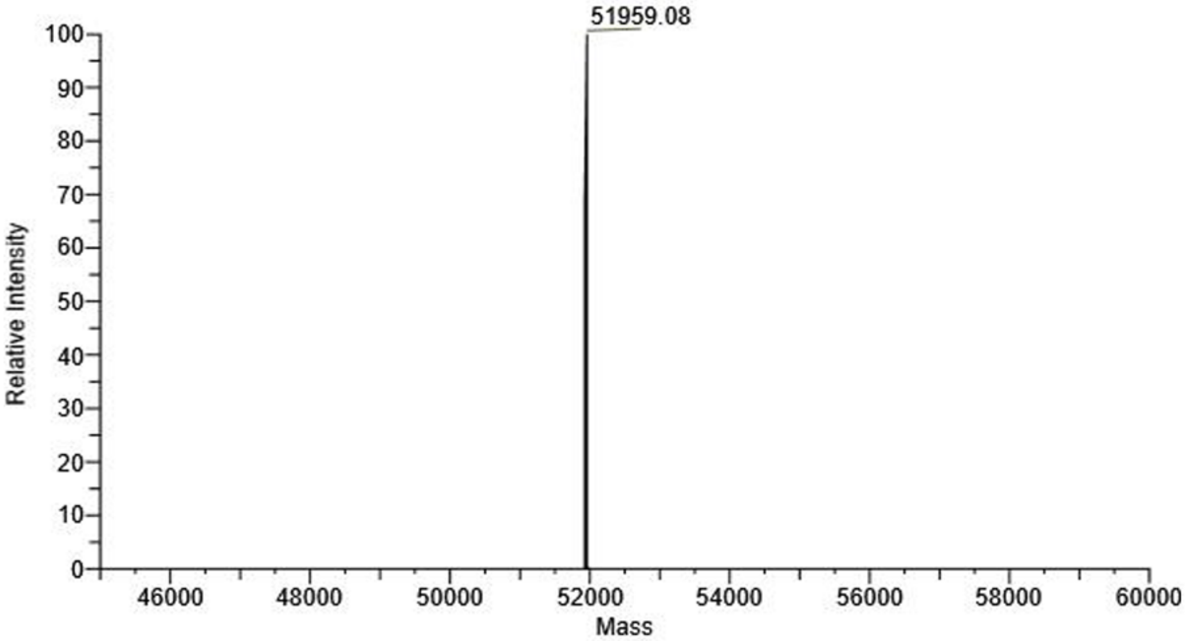


**Fig.S12 Mass Spectroscopy of OmcZ$^+$**. Reported mass of 51,959Da. Theoretical weight of OmcZ$^+$ with 8 hemes and strepII tag is 51,905Da.

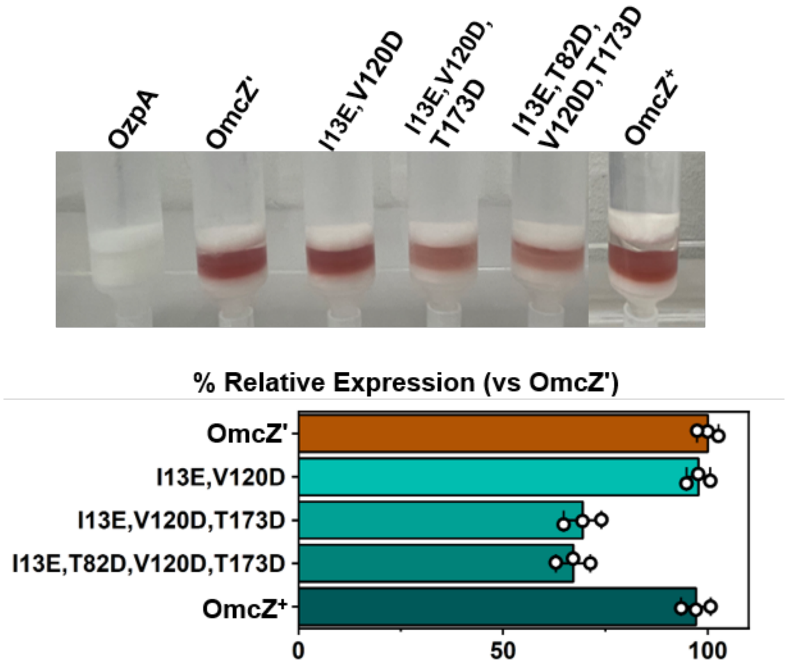


**Fig.S13 Relative OmcZ'$_{50}$ strepII expression**. Supernatant from 1L cultures of OzpA and OmcZ' variants. 500uL of high capacity strepII resin as shown binds ~15.5mg of protein. Color saturation determined using imageJ with OzpA as control.

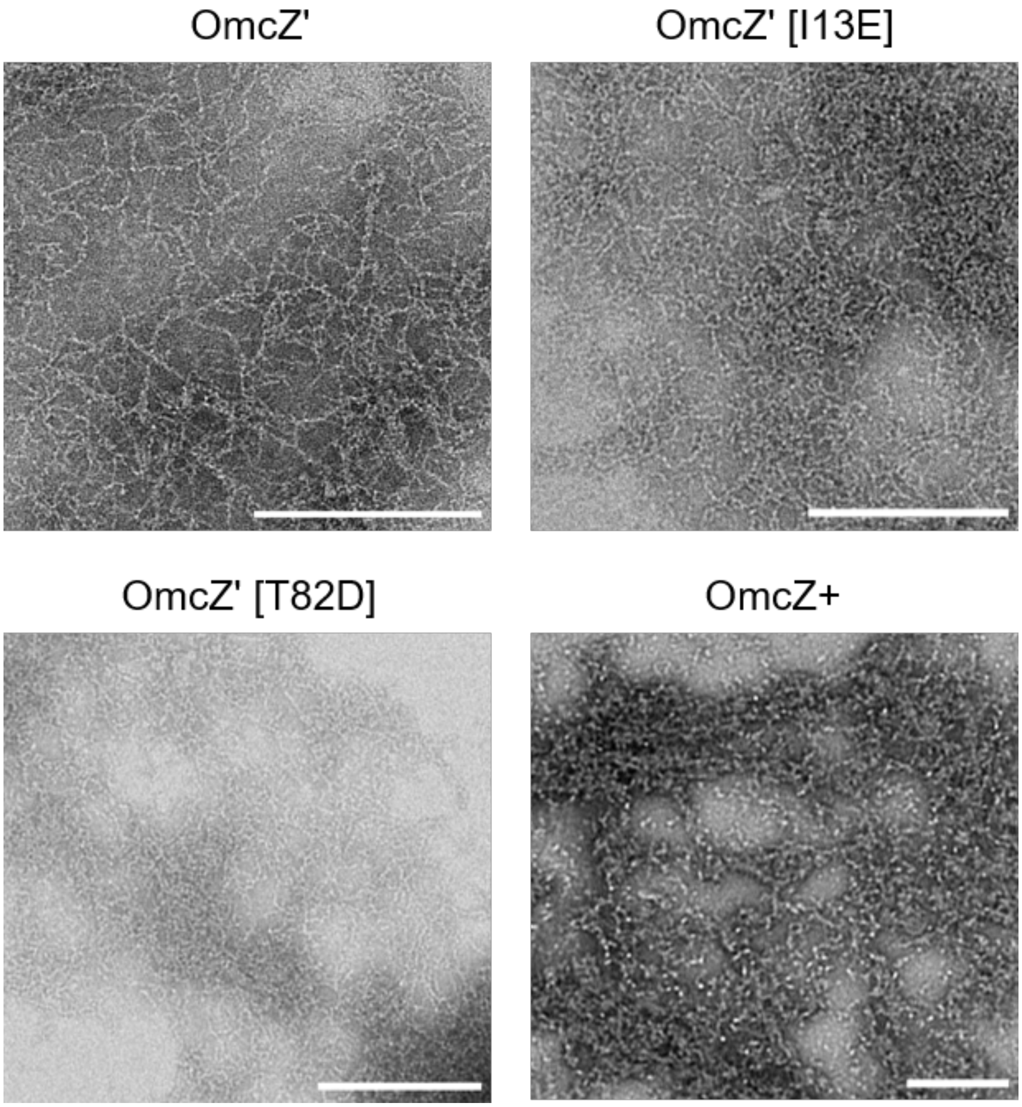


**Fig.S14 TEM of OmcZ' and variants.** Samples resuspended in 20mM HEPES pH8 and stained with 1% PTA.

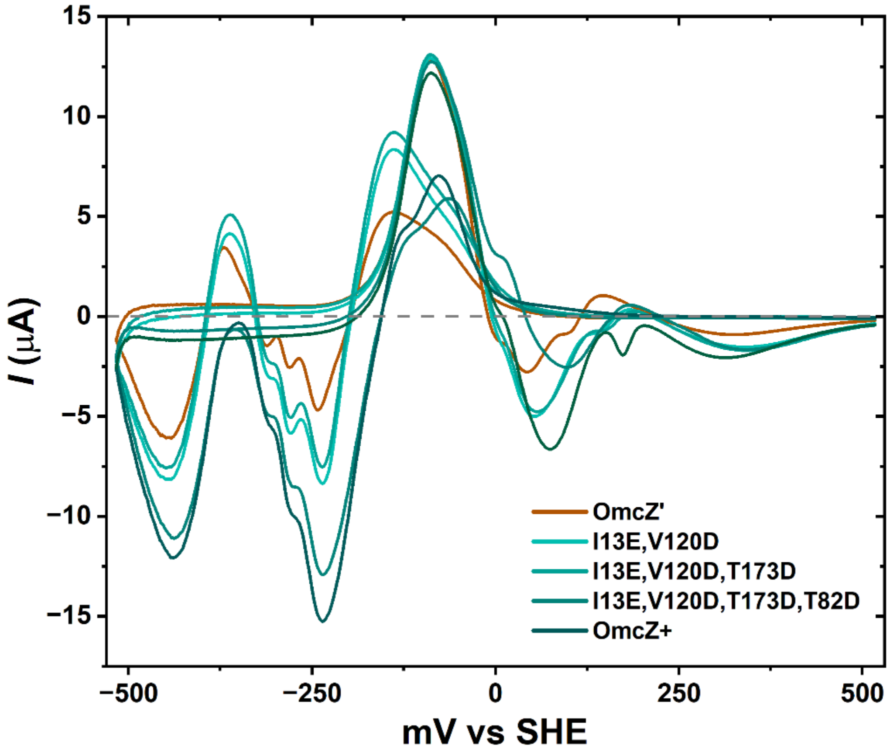


**Fig.S15 Cyclic voltammetry of purified OmcZ' and engineered nanowires.** Samples resuspended in 0.1X PBS at pH7.4 to 40μM on gold screen printed electrodes with Ag reference. Data from initial cathodic → anodic sweep measured at 50mV/s scan rate after background subtraction.

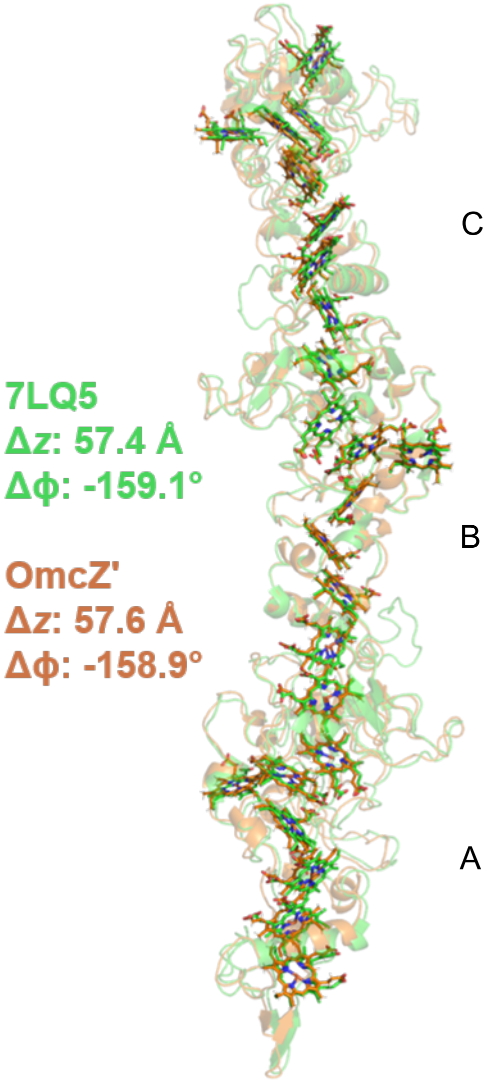


**Fig.S16 Native OmcZ alignment with OmcZ'.** Alignment (by chain B) of native *Geobacter sulfurreducens* OmcZ (PDB: 7LQ5, green) with *Shewanella oneidensis* OmcZ' (orange)

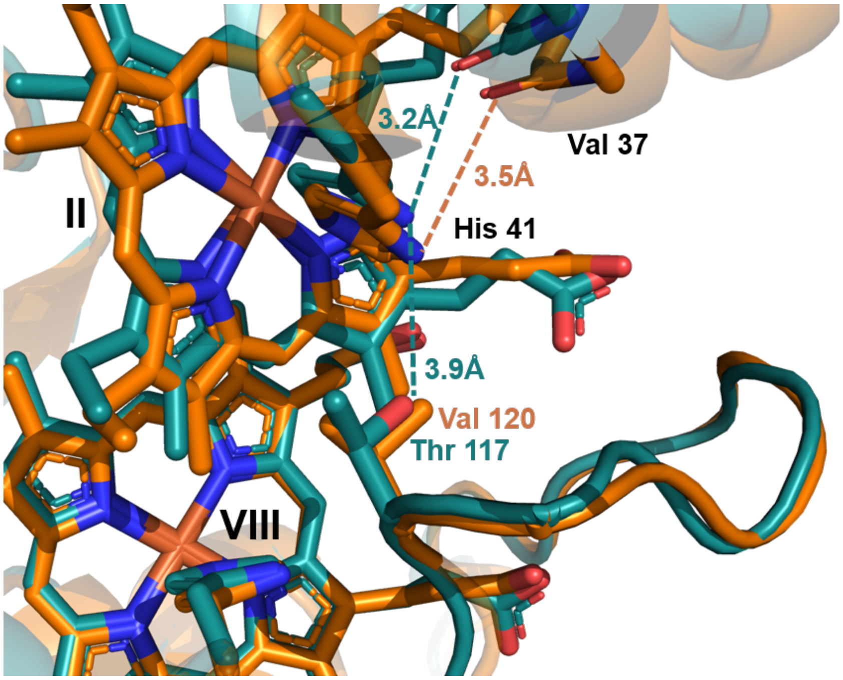


**Fig.S17** OmcZ' (orange) and OmcZ$^{+}$ (teal) overlay of engineered mutations S119D and V120T near heme VIII-II pocket

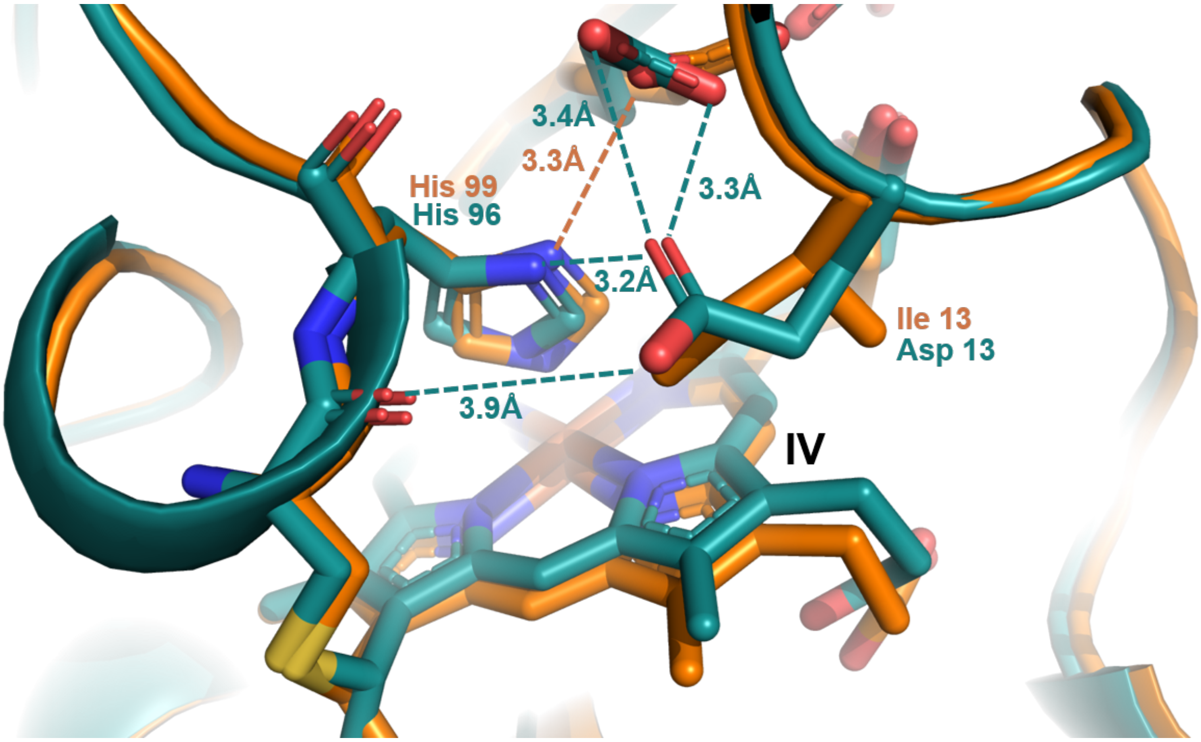


**Fig.S18** OmcZ' (orange) and OmcZ$^+$ (teal) overlay of engineered mutation I13D near heme IV pocket

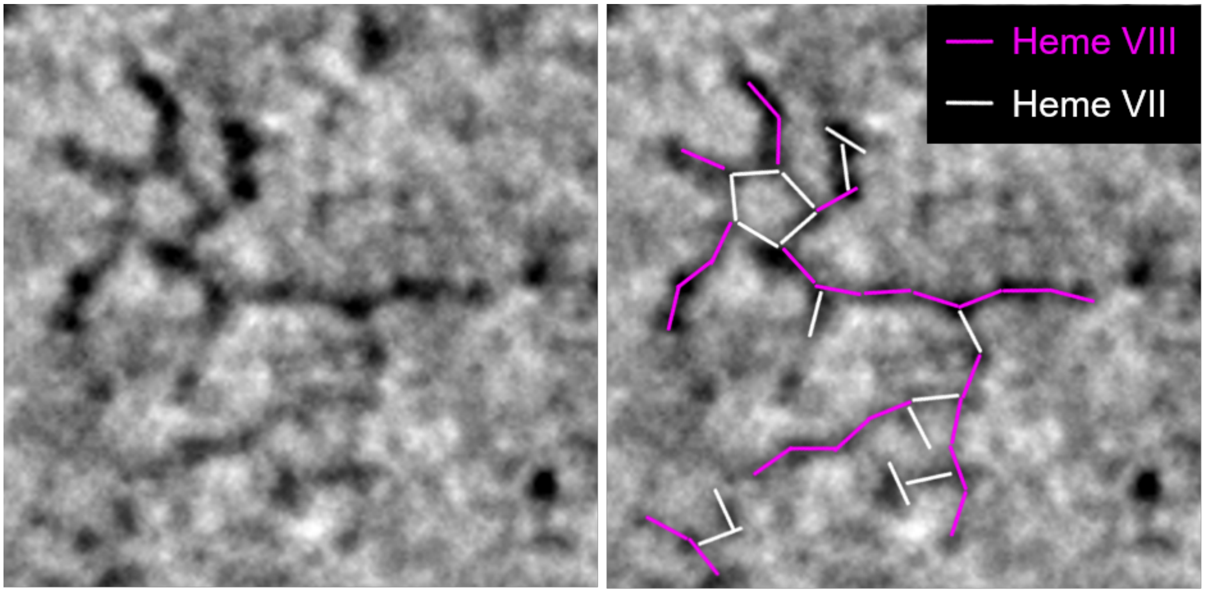


**Fig.S19 Rough estimation of Heme VIII vs Heme VII nanowire assembly in OmcZ⁺.** Representative methodological sample of direct inspection used in inter-subunit assembly (magenta: HemeVIII / white: HemeVII) micrograph inference.

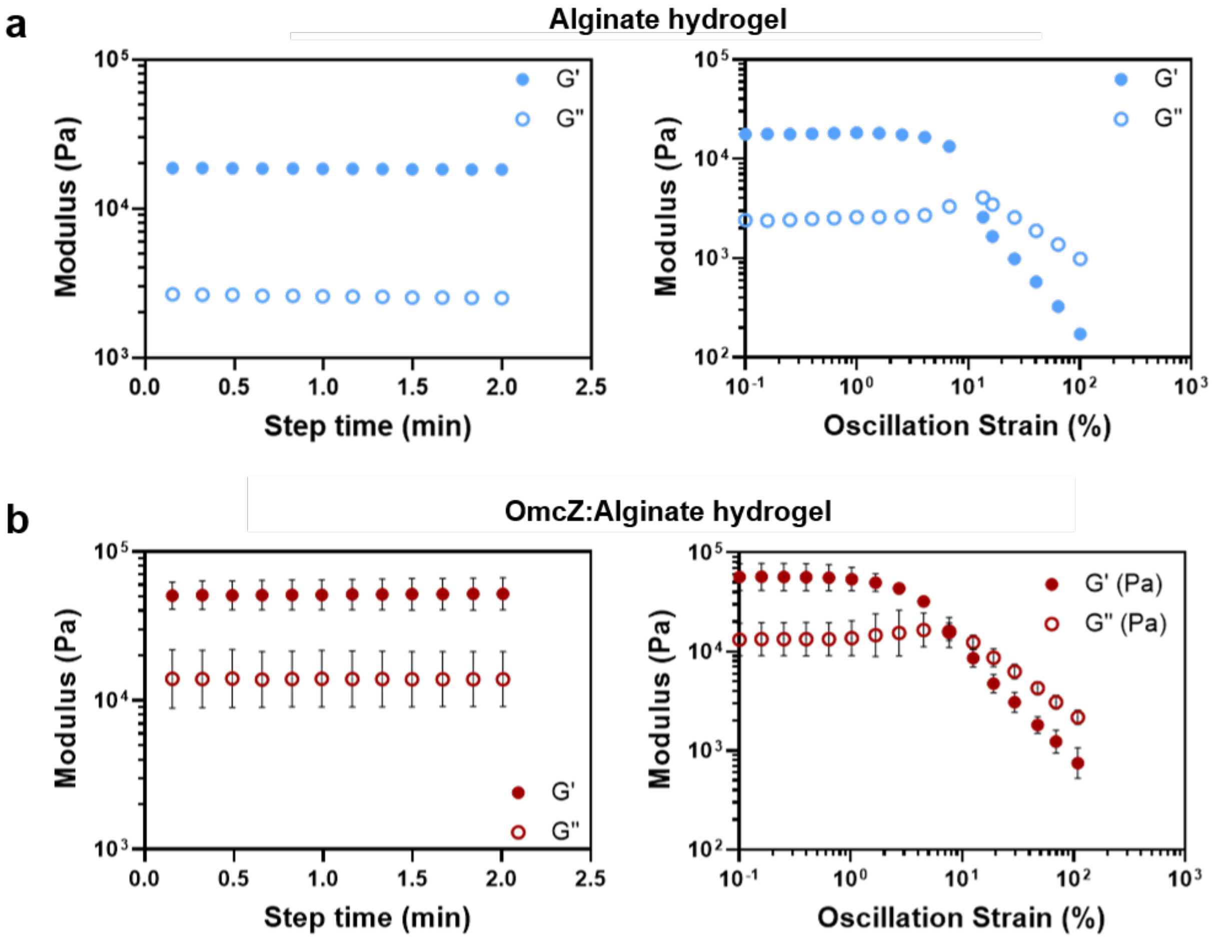


**Fig.S20 Alginate hydrogel rheometry.** (a) Alginate hydrogel. (b) Hybrid OmcZ'::alginate hydrogel

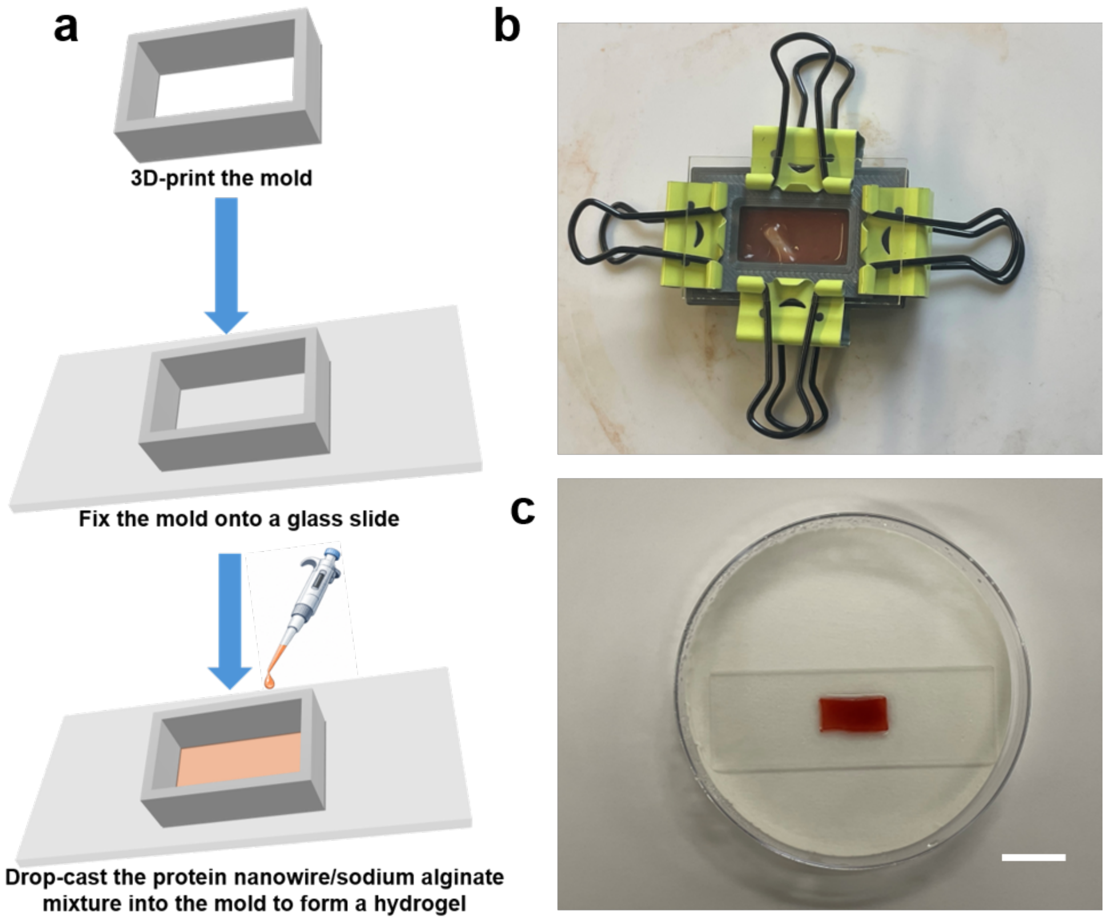


**Fig.S21 Fabrication of the hybrid OmcZ'::alginate hydrogel**. (a) Schematic showing the fabrication process of the hybrid OmcZ'::alginate hydrogel. (b) Photograph of the actual experimental setup. (c) Photograph of the fabricated hybrid OmcZ'::alginate hydrogel. Scale bar, 2 cm.

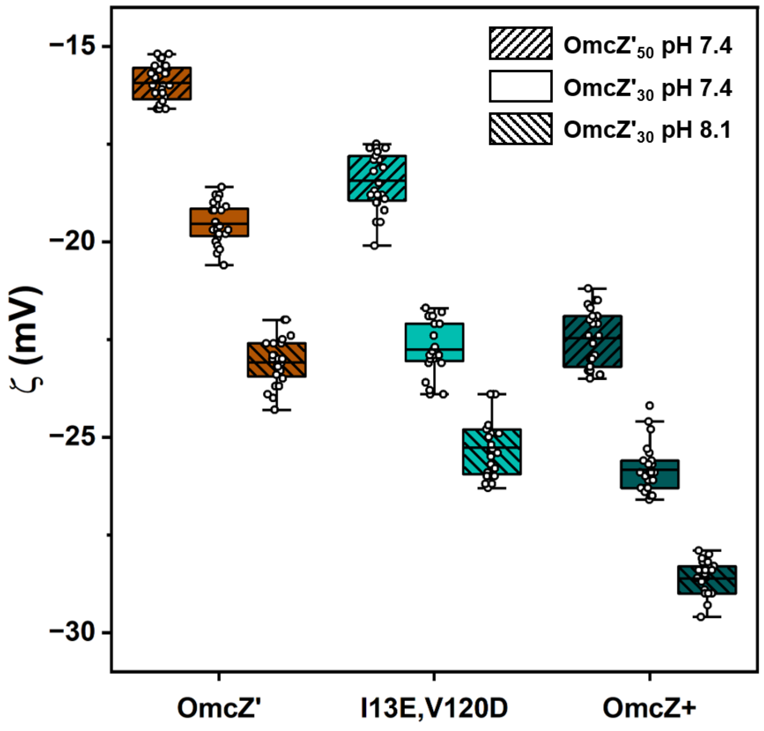


**Fig.S22 Zeta potential of OmcZ' variants**. Pro-peptide (OmcZ'$_{50}$) or nanowire (OmcZ'$_{30}$) zeta potential at varying pH.

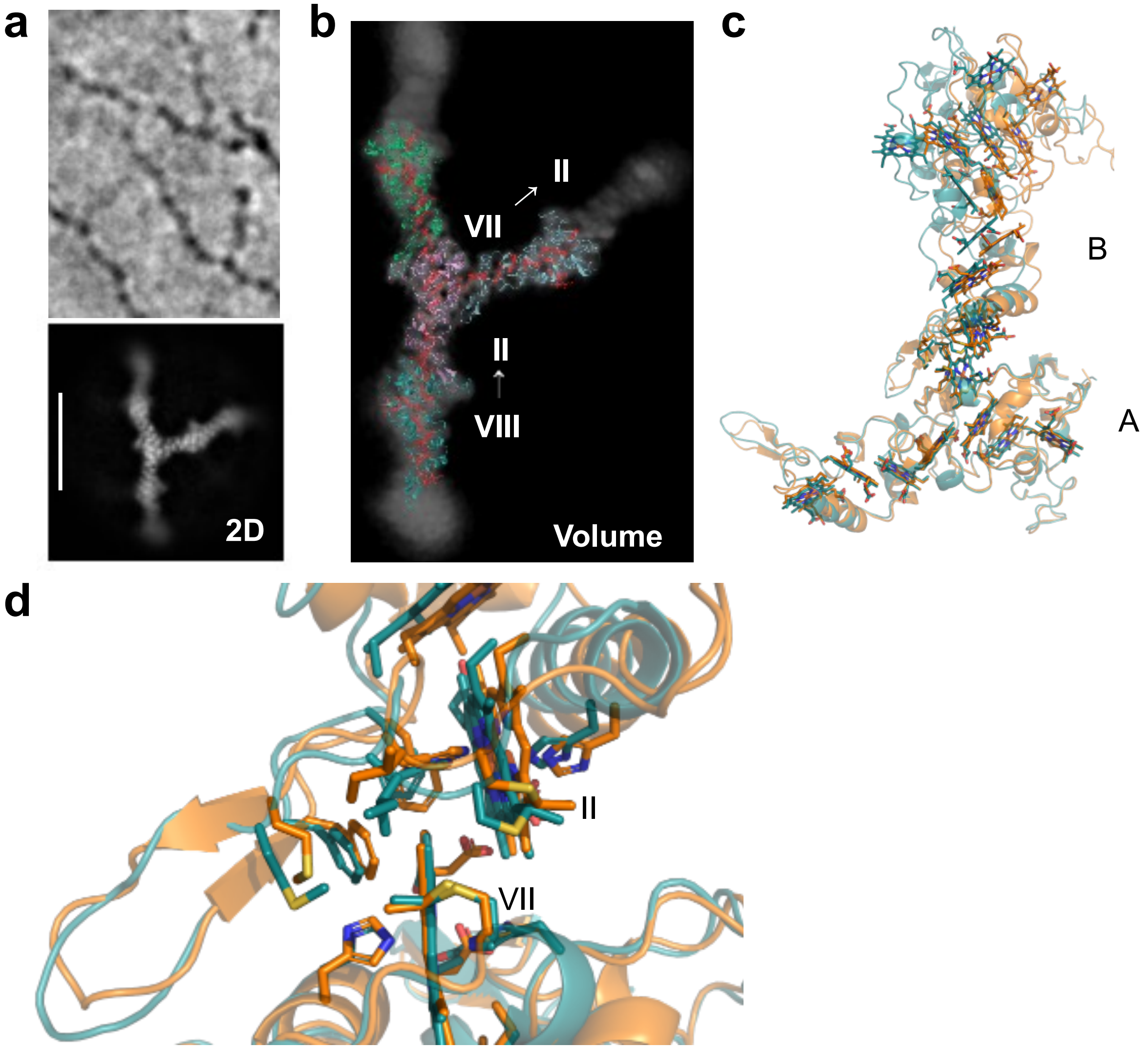


**Fig.S23 OmcZ' Heme VII-II assembly**. (a) Top, representative micrograph slice from OmcZ'. Bottom, sample 2D class average. (b) Low-resolution map with rough density fitting of OmcZ' nanowire (36HZ) and single subunit (chain B) at HemeVII. (c) OmcZ' HemeVII-II dimer (orange) aligned with OmcZ$^{+}$ HemeVII-II dimer (teal) by chain A. (d) Alignment from (c) at HemeVII and Heme II interface.
